# Transplanting ANXA1⁻ CD8⁺ Naïve T cells Delay Aging Through Senolysis

**DOI:** 10.64898/2026.02.21.707223

**Authors:** Yue Wu, Shang Guo, Fengjiao Zhang, Fangzhou Lou, Yan Li, Yang Sun, Qingqing Shen, Liuqi Zhao, Xiaojie Cai, Zhikai Wang, Qing Yang, Xichen Zheng, Min Gao, Xiangxiao Li, Siyu Deng, Zhenyao Xu, Honglin Wang

## Abstract

Immunosenescence is a hallmark of aging, yet strategies using defined immune subsets to counteract it are largely unexplored. We performed paired scRNA/TCR-seq on CD45⁺ cells from human bone marrow (15 donors, 3-91 years). T cells were the most altered lineage, with CD8⁺ naïve T cell contraction and functional impairment. We identified an expanding ANXA1⁺ CD8⁺ naïve subset with senescence signatures and impaired function. ANXA1-deficient CD8⁺ T cells exhibited increased resting stemness and enhanced activation/cytotoxicity upon stimulation. ANXA1⁻ cells showed senolytic activity *in vitro* and reduced senescence burden *in vivo*. Monthly transfer of syngeneic ANXA1⁻ CD8⁺ naïve T cells into aged mice extended median lifespan by >30 weeks, improving cardiac function, bone density, motor coordination, and marrow immune microenvironment. Our study identifies ANXA1 as critical in CD8⁺ naïve T cell aging and establishes ANXA1⁻ cell transfer as a strategy to counter immunosenescence and promote healthy aging.

## INTRODUCTION

The progressive decline of the immune system, termed immunosenescence, is a fundamental driver of systemic aging, rendering the elderly vulnerable to infections, malignancies, and autoimmune diseases (*1, 2*). A central hallmark of this process is the remodeling of the T cell compartment, historically characterized by the accumulation of highly differentiated effector and memory cells, often in response to persistent viral infections like cytomegalovirus (CMV) (*3–5*). These senescent or exhausted T cells contribute to a chronic, low-grade inflammatory state known as “inflammaging“(*6*), which can promote degenerative pathologies in distal tissues.

Immunosenescence involves intricate and broad alterations across immune cell subsets(*7–10*), lymphoid microenvironments(*11, 12*), and circulating factors(*9*). The dynamic interplay between innate and adaptive immunity is subject to immunosenescent changes at every level, including shifts in composition and functionality of T(*13*), B(*9*), natural killer (NK)(*14*) cells, and macrophages, directly impacting all processes including antigen-presenting cell (APC) activation(*15*), immune clearance(*16*) and further regulation of immune responses. Recent multi-omic atlases of human peripheral blood and thymus have confirmed that T cells are the most age-impacted lineage, undergoing robust transcriptional reprogramming and compositional shifts(*17, 18*). While the quantitative decline of the naïve T cell pool is a well-established feature of immunosenescence, a critical unanswered question is whether the remaining naïve T cells are mere bystanders or are themselves intrinsically compromised. The complexity and heterogeneity of age-related changes have so far impeded the identification of precise biomarkers that define such functionally distinct subsets within the naïve T cell pool. Crucially, these recent efforts have focused on peripheral blood or the thymus, while the human bone marrow—a foundational immunological niche that maintains a more stable, lifelong pool of immune cells — remains a critical, uncharacterized reservoir in this context(*18, 19*). Thus, a deeper, single-cell resolution atlas of a stable hematopoietic niche is required to dissolve this heterogeneity in immunosenescence in human.

Given the projected surge in aging-associated disorders with population aging, developing therapeutics to attenuate and potentially reverse functional deterioration is another critical pursuit(*20, 21*). It has been shown that aging, as well as immune-aging are both plastic processes that can be modified by habitual and pharmacological interventions(*20, 22, 23*). Therapeutic strategies against aging have shown promise, ranging from pharmacological senolytics that clear senescent cells to systemic rejuvenation. For example, targeting the SASP and resistance to apoptosis in senescent cells, 46 compounds that target senescent cell antiapoptotic pathways (SCAPs) were identified as potentially senolytic(*24–26*). Intermittent administration of the natural flavonoids procyanidin C1 inhibits SASP formation and selectively kills senescent cells(*27*). Meanwhile, the well-established heterochronic parabiosis (HPB) model has provided compelling evidence that exposure to youthful systemic milieu rejuvenates age-related impairments(*28–30*). However, the specific cellular effectors within a youthful systemic milieu that mediate these benefits remain largely unknown. Given the immune system’s capacity for systemic surveillance and its role in clearing damaged cells, we hypothesized that a functionally competent subset of immune cells could serve as a potent “endogenous senolytic” agent.

Here, we provide a comprehensive single-cell atlas of the human bone marrow CD45^+^ immune cells compartment across the lifespan. We identify the reduction of CD8⁺ naïve T cells as the central event of immune aging and, critically, discover that this is coupled with the emergence of a functionally distinct, senescence-like subpopulation defined by the surface expression of Annexin A1 (ANXA1). We identify ANXA1 as an intrinsic factor linking CD8⁺ naïve T cell aging to functional decline. Finally, we demonstrate that the targeted immunomodulation with adoptive transfer of ANXA1⁻ CD8⁺ naïve T cells serves as a powerful senolytic therapy, systemically ameliorating age-related disorders, extending both healthspan and lifespan in mice.

## RESULTS

### Single-cell atlas of the human bone marrow immune compartment with age highlights T cell population as a major site of transcriptional alterations

To capture the foundational changes of immunosenescence, we focused on the human bone marrow, a primary hematopoietic niche that maintains more stable and diverse of immune cell populations compared to the dynamic fluctuating peripheral circulation. We performed paired single-cell RNA sequencing (scRNA-seq) and T cell receptor sequencing (scTCR-seq) on CD45⁺ immune cells sorted from bone marrow aspirates of 15 healthy donors, who were stratified into five age groups spanning from childhood to old age (3 to 91 years) (**Fig. 1A, Fig. S1A** and **Table S1**). Successful scTCR-seq data was obtained from 13 of these donors. After quality control and filtering, a total of 107,587 cells were retained for downstream analysis. Unsupervised clustering based on the expression of signature marker genes identified 18 distinct immune cell populations (**Fig. S1B**). Major immune lineages, including T, B, NK, and myeloid cells, were visualized using t-Distributed Stochastic Neighbor Embedding (t-SNE), generating both an integrated projection and separate projections for each age group (**Fig. 1B**).

**Fig. 1.**
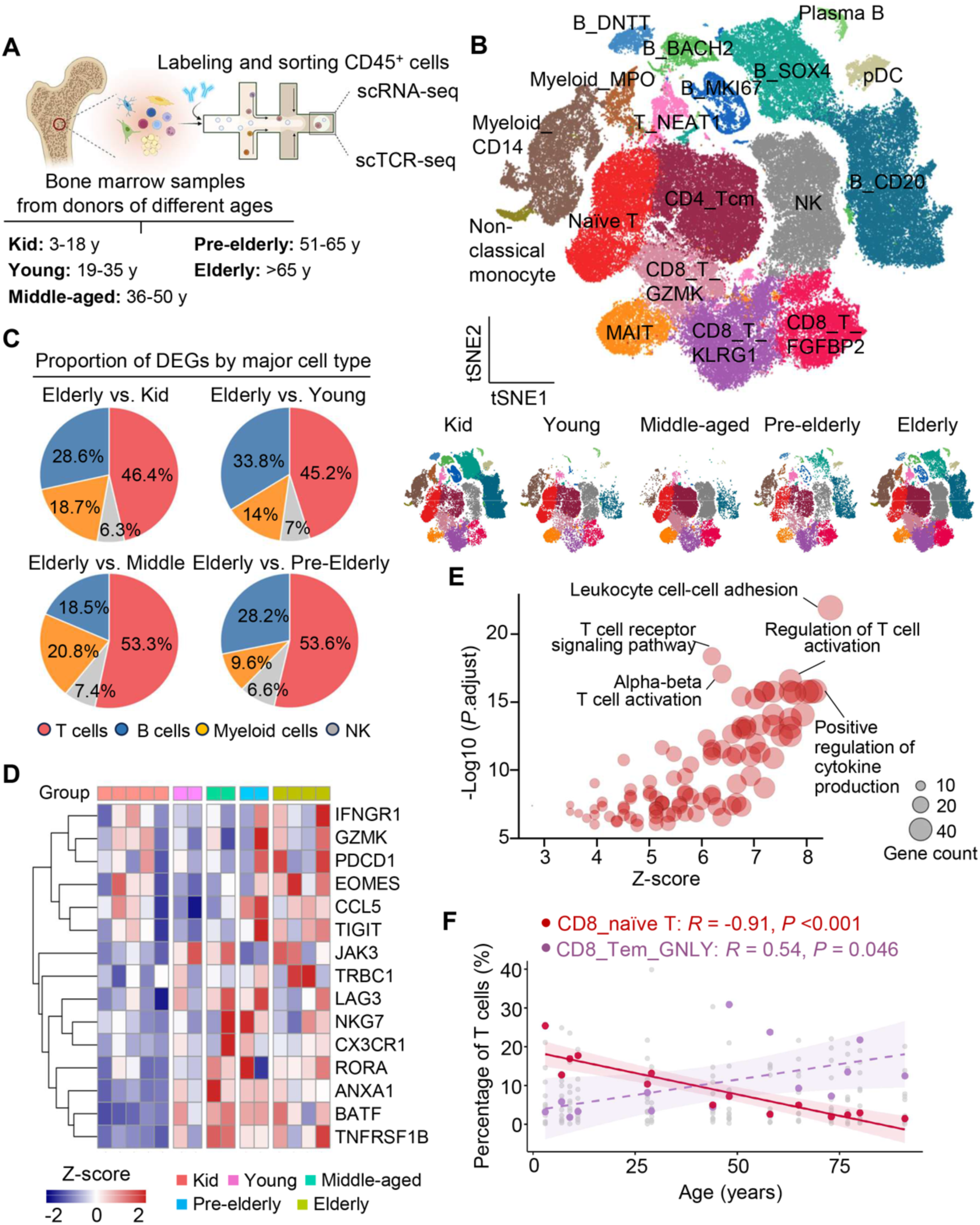
Single-cell characterization of the human bone marrow immune landscape across the lifespan reveals age-associated T cell alterations. **(A)** Schematic overview of the experimental design. CD45⁺ immune cells were isolated from bone marrow samples of 15 healthy donors aged 3 to 91 and subjected to paired single-cell RNA and TCR sequencing. Successful scTCR-seq data was obtained from 13 of these donors. Group definition will be referred as: Kid (3-18 y, n = 5 for scRNA-seq; n = 4 for scTCR-seq), Young (19-35 y, n = 2 for both scRNA-seq and scTCR-seq), Middle-aged (36-50 y, n = 2 for both scRNA-seq and scTCR-seq), Pre-elderly (51-65 y, n = 2 for both scRNA-seq and scTCR-seq), Elderly (>65 y, n = 4 for scRNA-seq and n = 3 for scTCR-seq). **(B)** T-SNE dimensional reduction and sub-clustering of 107,587 CD45⁺ immune cells from human bone marrow donors, shown as an integrated dataset with cell type annotations (top). The bottom panels show the same t-SNE projection, with cells split by their respective age groups defined in (**A**). **(C)** Pie chart displaying the proportional distribution of differentially expressed genes (DEGs) identified between the Elderly and other age groups. Proportions are categorized by major immune cell types: T cells, B cells, NK cells, and Myeloid cells. **(D)** Heatmap of selected DEGs in T cells across all age groups. The genes shown were identified as consistently dysregulated in the Elderly group when compared to all other age groups, derived from (**C**) Each row is scaled using a Z-score to visualize relative expression. **(E)** GO enrichment analysis of DEGs from Elderly group when compared to all other age groups. The bubble plot shows representative enriched pathways. **(F)** Correlation scatter plot of the proportions of key T cell subsets with donor ages in scRNA-seq data (n = 15). T cell subsets exhibiting a statistically significant correlation with age (*P* < 0.05) are highlighted in color, while all other subsets are depicted in grey. Pearson correlation coefficient (R) and *P*-values are indicated.

To investigate the transcriptional alterations that occur during aging, we performed a differential gene expression analysis comparing the Elderly group against all other age groups. When these differentially expressed genes (DEGs) were categorized by their cell lineage of origin, we found that T cells accounted for the largest proportion, suggesting T cells as the most transcriptionally dynamic during aging (**Fig. 1C**). We next examined genes consistently dysregulated in the Elderly group compared to all other age groups within T cells. A heatmap visualizing the expression patterns of selected genes across all donors, highlighting the age-associated upregulation of key genes relevant to T cell function, including *CCL5*, *NKG7*, *ANXA1*, and *TIGIT* (**Fig. 1D**). Furthermore, we performed gene ontology (GO) enrichment analysis on DEGs identified between Elderly and Kid donors across all cell types. This analysis also revealed significant enrichment of pathways such as “T cell receptor signaling pathway” and “positive regulation of cytokine production” (**Fig. 1F**), indicating a shift towards a dysregulated, pro-inflammatory state within the aged T cell compartment.

### Age-associated T cell remodeling is characterized by naïve T cell contraction, reduced TCR diversity, and altered functional capacity

Given the primary T cell alterations observed in the overall immune atlas, we performed a detailed sub-clustering and analysis focused on the T cell compartment (**Fig. S2A, B**). Compositional analysis revealed age-associated shifts across various T cell subsets (**Fig. S2C**), with a consistent reduction in the proportion of CD8⁺ naïve T cells. To quantify these changes, we correlated the abundance of each T cell subset with donor age. This analysis confirmed the contraction of CD8⁺ naïve T cells as the most statistically significant age-related alteration within the T cell lineage (R = −0.91, *P* < 0.001; **Fig. 1F, Fig. S2D**), accompanied by a corresponding expansion of terminally differentiated CD8⁺ effector memory T cell cluster (CD8⁺ Tem_GNLY) (**Fig. 1F**).

Beyond this quantitative decline observed in scRNA-seq, we next examined age-related alterations in the T cell receptor (TCR) repertoire diversity, a critical factor of immunity against novel pathogens(*31, 32*). Analysis of paired scTCR-seq data revealed reduction in repertoire diversity with increasing age. The highly diverse repertoire characteristic of Kid and Young donors, dominated by unique (singletons) and small T cell clones, progressively shifted towards a highly constricted repertoire in Elderly individuals. This aged repertoire was characterized by the predominance of a few large and hyperexpanded oligoclonal populations (**Fig. 2A**, **Fig. S3A**). This loss of diversity was quantified by the decrease in both the percentage of unique clonotypes (**Fig. 2B**) and the Inverse Simpson Index, a widely used diversity metric (**Fig. 2C**). Notably, these hyperexpanded clonotypes were predominantly found within the he CD8⁺ effector and memory T cell subsets (**Fig. 2D, Fig. S3B, D**). Chord diagram analysis further illustrated the shift of the clonal architecture, particularly within the CD8⁺ T cells across the age groups (**Fig. S3C**).

**Fig. 2.**
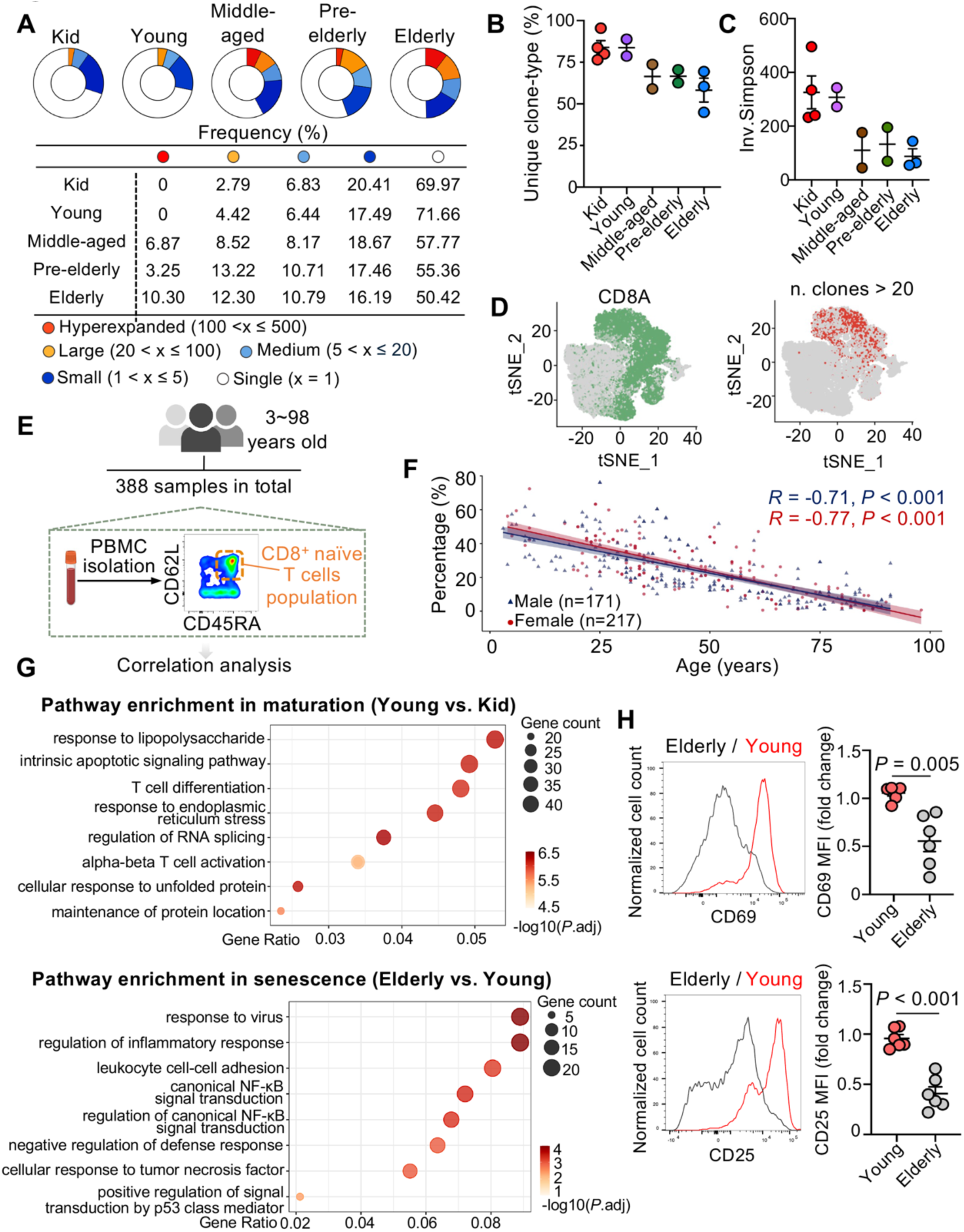
Age-associated remodeling of the T cell compartment is characterized by contraction of the naïve T cell pool, loss of TCR repertoire diversity, and functional decline. **(A)** TCR clonotype size distribution across age groups. Pie charts and a summary table quantify the frequency of clonotypes, categorized by size: Hyperexpanded (100 <x ≤ 500), Large (20 <x ≤ 100), Medium (5 < x ≤ 20), Small (1 < x ≤ 5), Single (x = 1). Data are shown for donors in the Kid (n = 4), Young (n = 2), Middle-aged (n = 2), Pre-Elderly (n = 2), and Elderly (n = 3) groups. **(B)** Quantification of unique TCR clonotypes (singletons) across age groups. The plot displays the percentage of singletons within the total T cell repertoire for each donor in the scTCR-seq cohort. **(C)** TCR repertoire diversity across age groups, measured by the Inverse Simpson Index. Each point represents the diversity score for an individual donor, where a higher index value indicates greater repertoire diversity. **(D)** T-SNE projection of T cells, with cells colored by their normalized CD8A expression level (left). The same plot is shown on the right, highlighting the location of large and hyperexpanded clonotypes. **(E)** Schematic of the large-cohort validation study and representative flow cytometry gating for CD8⁺ naïve T cells (CD45RA⁺ CD62L⁺) in human PBMCs. **(F)** Correlation scatter plot of the percentage of CD8⁺ naïve T cells with age in a large cohort of 388 healthy donors (217 males, 171 females). Pearson correlation coefficient (R) and *P*-values are indicated. **(G)** Comparative GO enrichment analysis for DEGs in naïve T cells. Bubble plots display pathways enriched during ‘Maturation’ (DEGs from Young vs. Kid comparison, top) and ‘Senescence’ (DEGs from Elderly vs. Young comparison, bottom). Bubble size is proportional to the gene count, and color intensity corresponds to the adjusted P-value. **(H)** Representative histograms and quantification of CD69 and CD25 expression on CD8⁺ naïve T cells from Young (n = 6) and Elderly (n = 6) donors after 12 hours of stimulation with anti-CD3/CD28 antibodies. MFI, mean fluorescence intensity. Data are presented as mean ± SEM. *P*-values were determined by Pearson correlation analysis and two-tailed unpaired t-test.

To determine whether the findings from our bone marrow sequencing analysis extend to the peripheral circulation, we quantified the proportion of CD8⁺ naïve T cells (defined as CD45RA⁺CD62L⁺) in peripheral blood mononuclear cells (PBMCs) from a large cohort of 388 healthy donors aged 3 to 98 years using flow cytometry (**Fig. 2E**). This large-scale flow cytometry analysis confirmed a significant and progressive decline in the frequency of circulating CD8⁺ naïve T cells with age, a trend observed in both male and female donors (**Fig. 2F**). When stratified into age brackets, this decline was clear throughout the lifespan (**Fig. S2E**). This reduction in peripheral CD8⁺ naïve T cells mirrored the trend observed in our bone marrow scRNA-seq cohort, indicating that the loss of naïve T cells is a systemic characteristic of human aging.

Beyond the observed decline in frequency, we next investigated whether the functional capacity of CD8⁺ naïve T cells is altered during aging, distinct from the changes occurring during early-life maturation. GO enrichment analysis was performed on DEGs of CD8⁺ naïve T cells identified from two comparisons: Young versus Kid donors (representing immune maturation) and Elderly versus Young donors (representing aging). This comparative analysis revealed that while pathways related to T cell activation and development were prominent during maturation (Young vs. Kid), pathways associated with inflammatory responses were enriched specifically during the aging phase (Elderly vs. Young) (**Fig. 2G**). Functionally, we isolated CD8⁺ naïve T cells from young and elderly donors and assessed their activation potential upon TCR stimulation. Naïve T cells from elderly donors exhibited lower upregulation of both the early activation marker CD69 and the IL-2 receptor alpha chain CD25 (**Fig. 2H**), indicating a compromised activation response in aged CD8⁺ naïve T cells.

### ANXA1 surface expression identifies an age-expanding CD8⁺ naïve T cell subset with senescence-associated features

The observed functional hypo-responsiveness of aged CD8⁺ naïve T cells prompted us to explore the underlying molecular basis for this decline. Gene Set Enrichment Analysis (GSEA) comparing the transcriptomes of CD8⁺ naïve T cells from elderly versus young donors revealed enrichment of the Hallmark “Cellular Senescence” and “Senescence-Associated Secretory Phenotype (SASP)” gene sets in aged cells, indicating the activation of a senescence-like program during T cell aging (**Fig. 3A**). We further examined the expression patterns of age-associated gene signatures across all donor. This analysis revealed a coordinated module of genes upregulated with age, with *ANXA1* (encoding Annexin A1) emerging as a key candidate alongside known pro-inflammatory genes such as *CCL5* (**Fig. 3B, C**). However, flow cytometric validation showed that while CCL5 protein expression was detectable, it did not have a distinct correlation in CD8⁺ naïve T cells with age (**Fig. S4A**).

**Fig. 3.**
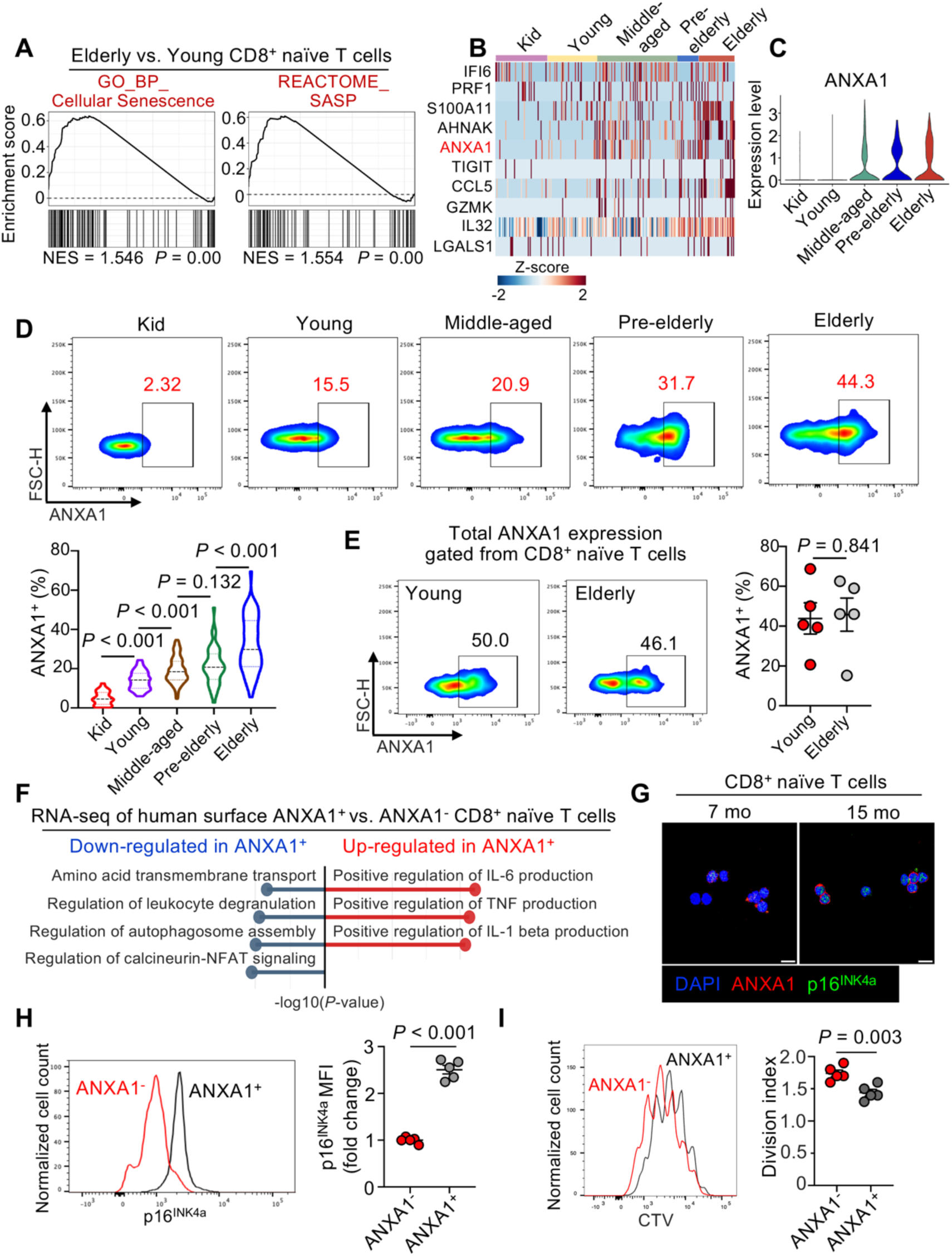
ANXA1 expression identifies an age-expanding CD8⁺ naïve T cell subset with features of cellular senescence. **(A)** Gene Set Enrichment Analysis (GSEA) comparing CD8⁺ naïve T cells from Elderly versus other groups of donors. Plots show significant enrichment of the Cellular Senescence (GO:0090398) and Senescence-Associated Secretory Phenotype (SASP; R-HSA-2559582) gene sets in the Elderly group. NES, normalized enrichment score. **(B)** Heatmap displaying the expression of senescence-associated signature genes in CD8⁺ naïve T cells across all donors. Each row is scaled using a Z-score. **(C)** Violin plot showing the normalized expression level of ANXA1 in CD8⁺ naïve T cells across the five age groups from scRNA-seq data. **(D)** Quantification of ANXA1 surface expression in CD8⁺ naïve T cells across the five age groups, as measured by flow cytometry. Each point represents an individual donor (Kid, n = 31; Young, n = 98; Middle-aged, n = 93; Pre-Elderly, n = 52; Elderly, n = 63). **(E)** Representative flowcytometry plots and summary quantification of total ANXA1 expression levels in CD8⁺ naïve T cells from human PBMCs (n = 5 per group). **(F)** Pathway enrichment analysis comparing human ANXA1⁺ versus ANXA1⁻ CD8⁺ naïve T cells from RNA-seq. The bubble plot shows key pathways significantly altered between the two subsets. Bubble size corresponds to the number of genes, and color indicates enrichment significance. **(G)** Representative immunofluorescence images for ANXA1 and p16^INK4a^ expression in human CD8⁺ naïve T cells. Scale bar, 10 µm. **(H)** Representative flow cytometry histogram and quantification p16^INK4a^ expression in human ANXA1⁺ versus ANXA1⁻ CD8⁺ naïve T cells (n = 5 per group). **(I)** Representative histogram of Cell Trace Violet (CTV) dye dilution in human ANXA1⁻ and ANXA1⁺ CD8⁺ naïve T cells after 72 hours of stimulation. Data are presented as mean ± SEM. *P*-value was determined by one-way ANOVA and two-tailed unpaired t-test.

ANXA1 is a pleiotropic calcium-dependent phospholipid-binding protein with diverse immunomodulatory roles(*33, 34*). While classically known for mediating the anti-inflammatory effects of glucocorticoids(*35*), primarily through actions on myeloid cells recent studies have established its involvement in T cell biology. For instance, it has been shown to enhance the inhibition function of Treg cells in triple-negative breast cancer(*36*) and potentially elevate pathogenic T cell responses in autoimmunity such as systemic lupus erythematosus (*37, 38*). Despite these roles in activated or pathological T cell states, the function of ANXA1 within the naïve T cells, particularly its expression dynamics and significance during the physiological aging process, has remained largely unexplored. Notably, ANXA1 can reside intracellularly but can also be externalized to the plasma membrane upon cellular activation or stress, where its surface expression may reflect distinct functional states(*39–41*).

To further investigate the heterogeneity within aged CD8⁺ naïve T cells, we performed sub-clustering on this population from our scRNA-seq data (**Fig. S4B**). This analysis identified six distinct subclusters (C1-C6), one of which, C4, was uniquely characterized by high expression of ANXA1 (**Fig. S4C**). Correlation analysis revealed that the relative abundance of this ANXA1⁺ (C4) subcluster significantly increased with donor age, exhibiting the strongest positive correlation among all identified subclusters (**Fig. S4D**). Importantly, both the ANXA1⁺ (C4) and ANXA1⁻ (C1-3, C5-6) subsets retained the classical naïve T cell gene signature, characterized by high expression of *TCF7* and *SELL* and low expression of canonical memory or effector markers like *EOMES* and *GZMB* (**Fig. S4E, F**).

Building on the identification of an age-expanding ANXA1⁺ CD8⁺ naïve T cell subset in bone marrow, we next investigated its presence and characteristics in peripheral blood by assessing ANXA1 protein localization and age-related expression dynamics. Analysis of our large human PBMC cohort demonstrated the progressive increase in the frequency of surface ANXA1-expressing CD8⁺ naïve T cells from childhood through old age (**Fig. 3D**). Notably, ANXA1 total expression of CD8⁺ naïve T cells didn’t show the same pattern with the donor age (**Fig. 3E**). Subsequently, this age-dependent expansion of an ANXA1⁺ CD8⁺ naïve T cell population with potential functional alterations was also conserved in mice, where surface expression of ANXA1 in both bone marrow and PBMCs was significantly elevated on CD8⁺ naïve T cells from aged compared to young animals (**Fig. S4G, H**).

The distinct surface expression of ANXA1 enabled the isolation of surface ANXA1⁺ and ANXA1⁻ CD8⁺ naïve T cells via fluorescence-activated cell sorting (FACS) for subsequent transcriptomic profiling. Transcriptomic analysis revealed that ANXA1⁺ CD8^+^ naïve T cells exhibited downregulation of key T cell activation pathways, including calcineurin–NFAT signaling, alongside enrichment of senescence-associated secretory phenotype (SASP), such as IL-6, TNF, and IL-1β (**Fig. 3F**). Consistent with this senescence-like transcriptional profile, sorted ANXA1⁺ CD8^+^ naïve T cells displayed increased protein expression of the canonical senescence marker p16^INK4a^ in both immunofluorescence (**Fig. 3G**) and flow cytometry (**Fig. 3H**) results. Furthermore, ANXA1⁺ CD8^+^ naïve T cells showed impaired proliferative capacity relative to ANXA1⁻ cells (**Fig. 3I**).

### ANXA1 deficiency potentiates the activation and cytotoxicity of CD8⁺ T cells

To investigate ANXA1 functions of the observed senescence-like phenotype, we utilized *Anxa1* knockout (*Anxa1*^KO^) mice. We first assessed whether ANXA1 deficiency influences the basal state of CD8⁺ naïve T cells. Transcriptomic profiling revealed that resting *Anxa1*^KO^ CD8⁺ naïve T cells exhibited the upregulation of gene sets associated with T cell homeostasis, adhesion/mobility, and stemness (**Fig. S5A, B**). Then, upon activation with anti-CD3/CD28 antibodies, *Anxa1*^KO^ CD8⁺ naïve T cells demonstrated upregulation of pathways related to immune activation and effector function (**Fig. 4A, B**). Consistent with this, *Anxa1*^KO^ CD8⁺ naïve T cells demonstrated higher and more rapid surface expression of the activation markers CD69 and CD25 (**Fig. 4C, D**). Functionally, activated *Anxa1*^KO^ CD8⁺ T cells produced more IFNγ (**Fig. 4E**) and demonstrated superior degranulation capacity, as indicated by increased surface mobilization of CD107a (**Fig. 4F**).

**Fig. 4.**
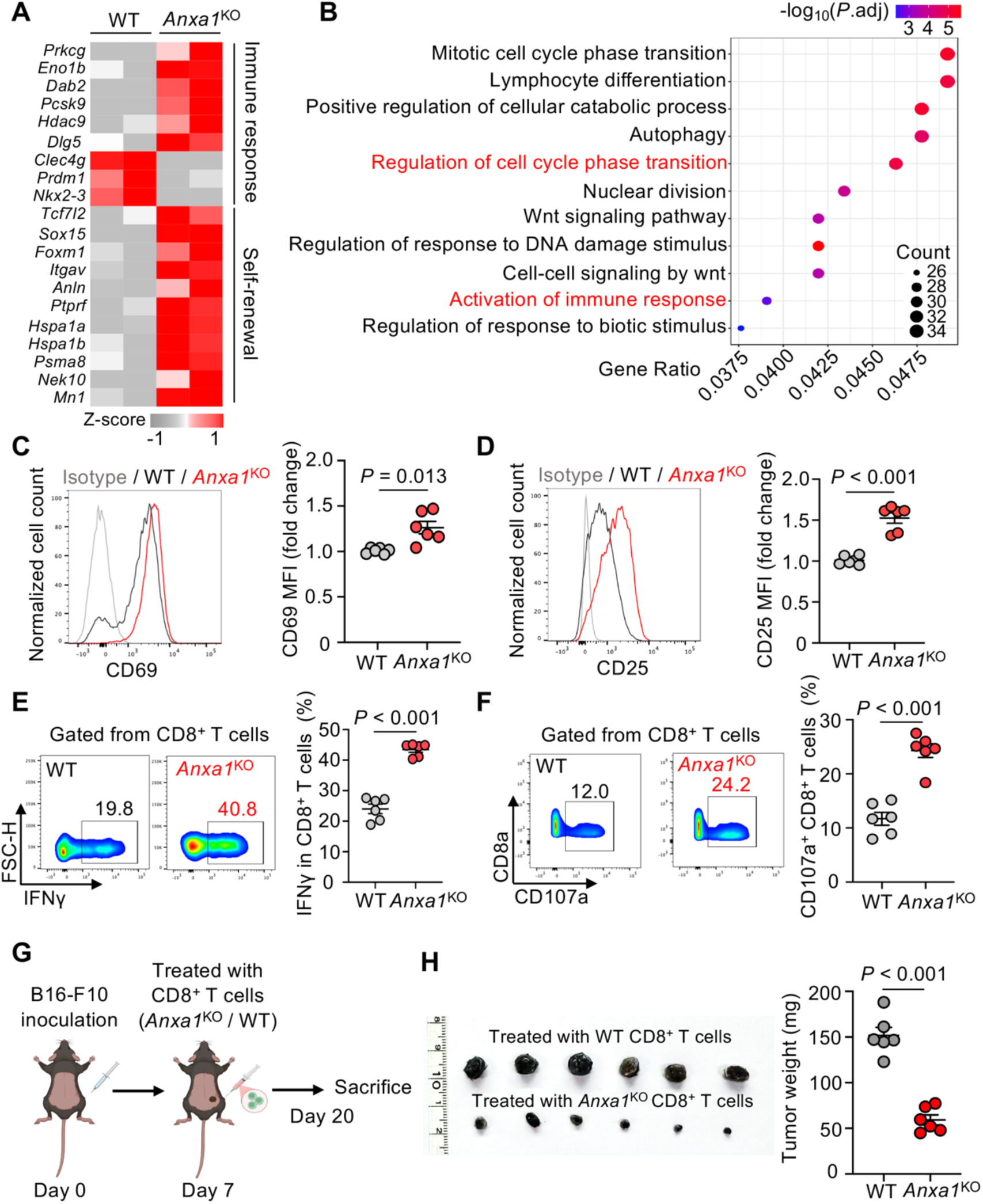
ANXA1 deficiency enhances the activation, cytotoxicity, and anti-tumor efficacy of CD8⁺ T cells. **(A)** Heatmap of DEGs in activated CD8⁺ naïve T cells from WT and *Anxa1*^KO^ mice. Cells were stimulated with anti-CD3/CD28 antibodies for 12 hours prior to RNA-seq. Each row is scaled using a Z-score. **(B)** GO enrichment of pathways upregulated in activated *Anxa1*^KO^ CD8⁺ naïve T cells compared with WT. (**C** and **D**) Representative flow cytometry histograms and quantification of activation markers CD69 (**C**) and CD25 (**D**) on WT and *Anxa1*^KO^ CD8⁺ naïve T cells 12 hours post-stimulation (n = 6 per group). **(E)** Representative flow cytometry plots and quantification of IFNγ-producing cells within WT and *Anxa1*^KO^ CD8⁺ T cells after stimulation (n = 6 per group). **(F)** Representative flow cytometry plots and quantification of the degranulation marker CD107a on WT and *Anxa1*^KO^ CD8⁺ T cells after stimulation (n = 6 per group). **(G)** Schematic of the B16-F10 inoculation model. C57BL/6 mice were inoculated subcutaneously with B16-F10 melanoma cells, followed by adoptive transfer of CD8⁺ T cells isolated from either WT or *Anxa1*^KO^ mice. **(H)** Representative image and quantification of excised tumors at day 20 post-inoculation. (n = 6 per group). Data in quantification plots are presented as mean ± SEM. P-values were determined by two-tailed unpaired t-test.

To test if this enhanced *in vitro* effector function translates to improved anti-tumor immunity *in vivo*, we employed a B16-F10 melanoma model (**Fig. 4G)**. C57BL/6 mice bearing established subcutaneous B16-F10 tumors received adoptive transfer of CD8⁺ T cells isolated from either WT or *Anxa1*^KO^ CD8⁺ T cells. Mice receiving *Anxa1*^KO^ CD8⁺ T cells exhibited suppression of tumor growth and reduction in final tumor mass compared to recipients of WT T cells (**Fig. 4H**). These results demonstrate that ANXA1 deficiency enhances CD8⁺ T cell activation and effector function *in vitro*, with improved control of tumor growth *in vivo*.

### ANXA1⁻ CD8⁺ naïve T cells ameliorate senescence via a direct senolytic mechanism

The accumulation of senescent cells contributes to tissue aging and age-related pathologies(*42*). While pharmacological senolytics can eliminate senescent cells, the potential of the immune system—particularly T cells—to mediate this clearance represents a compelling therapeutic strategy(*16, 43*). Given that ANXA1 marks an age-expanding, senescence-associated CD8⁺ naïve T cells, while its absence enhances cytotoxic potential, we explored whether the ANXA1⁻ subset contributes to immune surveillance against senescent cells.

To investigate this potential and disentangle the effects of ANXA1 status from donor age, we first compared the *in vitro* functional capacity of surface ANXA1⁻ CD8⁺ naïve T cells sorted from young (8-week-old) versus aged (13-month-old) mice. Upon stimulation, ANXA1⁻ cells from both young and aged donors exhibited comparable proliferation and similar upregulation for the activation markers CD69 and CD25 (**Fig. S6A**). The differentiation results into cytotoxic T lymphocytes (Tc1 cells) were also indistinguishable between young and old donors (**Fig. S6B**). This functional equivalence was further validated in a U2OS tumor cell invasion assay, where ANXA1⁻ T cells from both young and old mice, but not ANXA1⁺ cells, effectively suppressed U2OS cancer cell migration (**Fig. S6C**). These results suggest that the enhanced functional capacity observed in these CD8⁺ naïve T cells is primarily associated with the ANXA1⁻ phenotype itself, rather than the donor’s chronological age.

We next directly assessed the senolytic capacity of ANXA1⁻ CD8⁺ naïve T cells. To dynamically visualize T cell interactions with senescent targets in real-time, we engineered fibroblasts to over-express p16^INK4a^ with GFP (**Fig. S7A, supplementary video 1**). Live-cell imaging over 72 hours revealed that ANXA1⁻ CD8⁺ naïve T cells actively migrated towards p16^INK4a^-GFP⁺ senescent fibroblasts, instead of normal primary fibroblasts, and established stable, prolonged contacts, ultimately mediated target cell clearance (**Fig. 5A, Fig. S7B**). In contrast, ANXA1⁺ CD8⁺ naïve T cells derived from the same donors failed to efficiently engage senescent cells, displaying minimal persistent interaction (**Fig. S7B**).

**Fig. 5.**
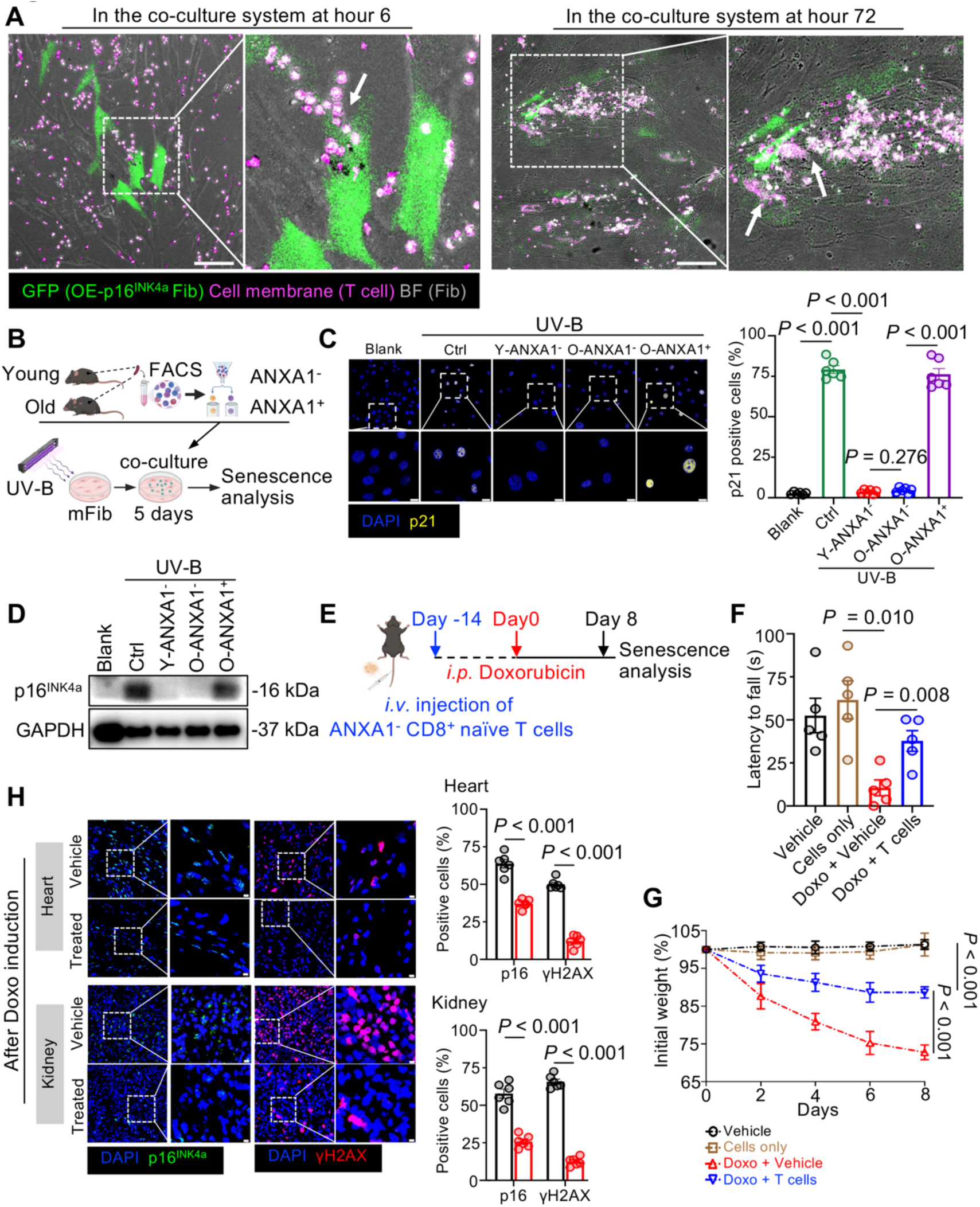
ANXA1⁻ CD8⁺ naïve T cells demonstrate senolytic activity *in vitro* and *in vivo*. **(A)** Representative stills from live-cell imaging of p16^INK4a^-GFP⁺ senescent fibroblasts (green), normal fibroblasts (grey) co-cultured with sorted surface ANXA1⁻ CD8⁺ naïve T cells (magenta), with arrows indicate interaction spots. **(B)** Schematic diagram of the in vitro UV-B induced senescence co-culture system. Mouse fibroblasts (mFib) were induced into senescence by UV-B radiation and then co-cultured for 5 days with different subsets of sorted CD8⁺ naïve T cells. **(C)** Representative immunofluorescence images and quantification of p21-positive fibroblasts after co-culture with the indicated T cell subsets (n = 6 per group). Scale bar, 10 µm. **(D)** Western blot analysis of p16^INK4a^ protein levels in senescent fibroblasts following co-culture. **(E)** Schematic of the doxorubicin (Doxo)-induced systemic senescence model. Mice received a single intravenous (*i.v.*) injection of ANXA1⁻ CD8⁺ naïve T cells prior to treatment with Doxo. **(F)** Rotarod test quantification showing motor coordination and endurance in mice (n = 5 per group). **(G)** Weight loss of mice in different groups (n = 5 per group) in Doxo-induced senescence model. Initial body weight was normalized to 100%. **(H)** Representative images and quantifications of p16^INK4a^ and γH2AX in the heart and kidney from Doxo-treated mice (n = 5 per group). Scale bar, 5 µm. Data in quantification plots are presented as mean ± SEM. P-values were determined by one-way ANOVA and two-tailed unpaired t-test.

To quantify this senolytic activity using established models, we employed UV-B irradiation to induce senescence in mouse fibroblasts *in vitro* (Fig. 5B). Consistent with previous data, only ANXA1⁻ CD8⁺ naïve T cells from both young (Y-ANXA1⁻) and old (O-ANXA1⁻) mice significantly reduced the number of p21-positive senescent fibroblasts, whereas ANXA1⁺ CD8⁺ naïve T cells from old mice (O-ANXA1⁺) were ineffective (**Fig. 5C**). This specific senolytic activity was further validated by reduced protein levels of p16^INK4a^ in senescent fibroblasts (**Fig. 5D**), with reduced expression levels of p53 and the DNA damage marker γH2AX (**Fig. S7C**). Consistent results were obtained using an independent Etoposide (ETOP)-induced senescence model (**Fig. S7D, E**). This senolytic activity was associated with the release of the cytotoxic effector molecule Granzyme B (GZMB) from ANXA1⁻ T cells upon engagement with senescent targets (**Fig. S7F**), suggesting the involvement of a granzyme-mediated cytotoxic mechanism.

We further employed a Doxorubicin (Doxo)-induced acute senescence model *in vivo* to assess the therapeutic potential of ANXA1⁻ CD8⁺ naïve T cells in systemic aging context (**Fig. 5D**). Doxo administration induced evident frailty in mice, including rapid weight loss and impaired motor coordination measured by the rotarod test (**Fig. 5F, G**). The infusion of ANXA1⁻ CD8⁺ naïve T cells administered prior to Doxo treatment significantly ameliorated these deficits, stabilizing body weight and restoring motor performance (**Fig. 5F, G**). This functional improvement correlated with reduced senescent cell burden *in vivo*. Immunofluorescence analysis of heart and kidney tissues at the study endpoint revealed that, mice that received ANXA1⁻ CD8⁺ naïve T cells had reduced level of p16^INK4a^- and γH2AX- positive senescence cells compared to control mice (**Fig. 5H**).

### Adoptive transfer of ANXA1⁻ CD8⁺ T cells specifically ameliorates age-related phenotypes

Based on the role of ANXA1 in defining an age-associated, functionally altered CD8⁺ naïve T cell subset and the enhanced potential of ANXA1-deficient T cells, we evaluated the therapeutic efficacy of adoptively transferring ANXA1⁻ CD8⁺ naïve T cells in naturally aged mice. To systematically assess this potential and determine the specificity of any observed effects, we conducted two complementary *in vivo* studies: a short-term (Day 250 endpoint) multi-arm trial comparing different transferred T cell populations (**Fig. 6A**), designed to dissect the cellular requirements for therapeutic benefit, and a long-term survival study focused on healthspan and lifespan (**Fig. 7A**).

**Fig. 6.**
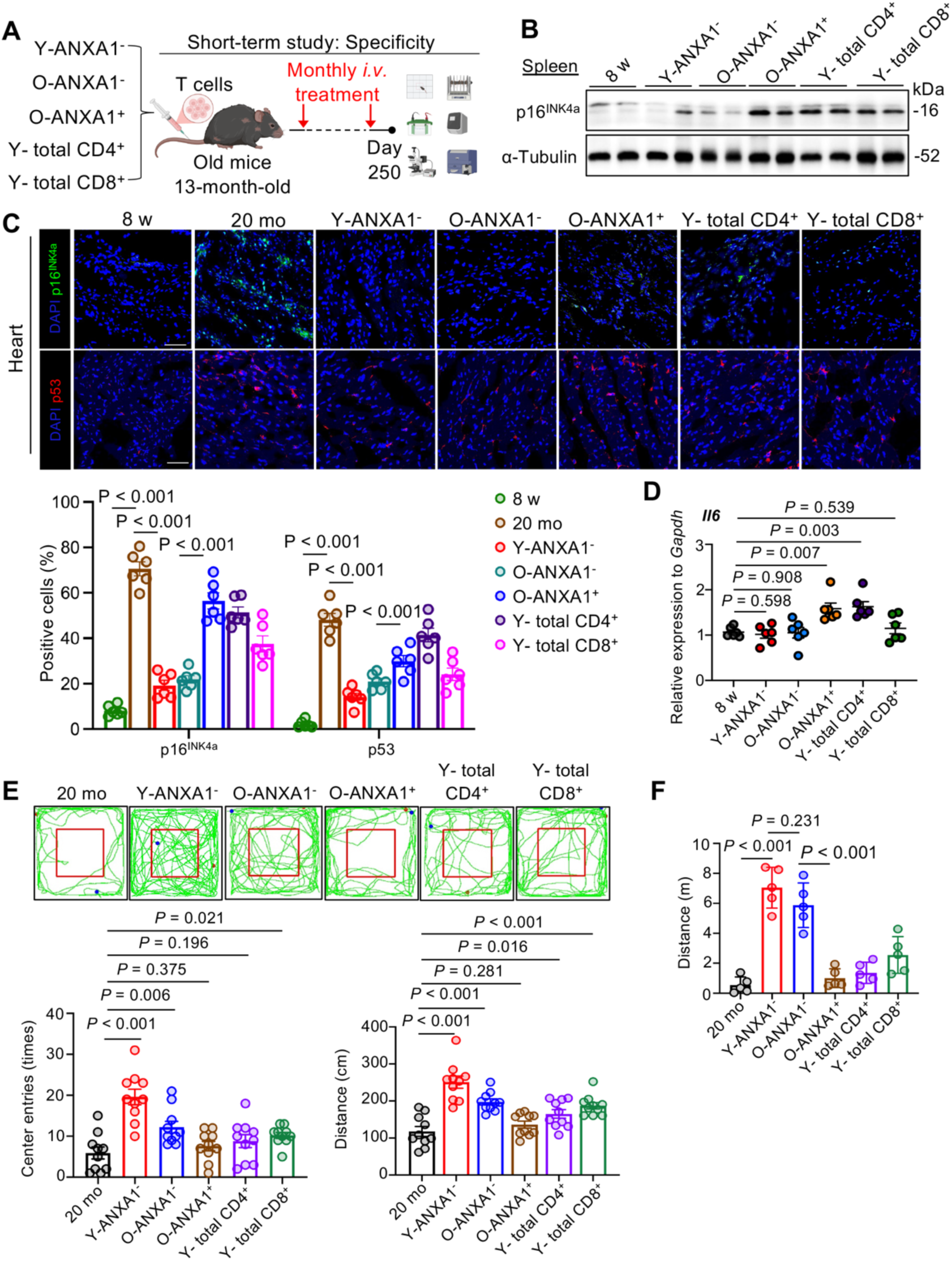
Short-term transfer of ANXA1⁻ CD8⁺ T cells specifically reduces senescence markers and improves physical function in aged mice. **(A)** Schematic of the short-term specificity study design. Old (13-month-old) mice received monthly intravenous (*i.v.*) transfer of one of five control T cell populations (specified in **B-F**) or vehicle, and were analyzed at Day 250 post-transfer. **(B)** Western blot for p16^INK4a^ expression in the spleen from the experimental groups (n = 5 per group). **(C)** Representative immunofluorescence images and quantifications of p16^INK4a^, p53 in the heart sections of different groups of mice (n = 6 per group). Scale bar, 50 µm. **(D)** RT-qPCR analysis of *Il6* mRNA expression in spleen tissue. Expression levels of the SASP factor *Il6* were quantified at Day 250 post-transfer in the indicated groups (n = 6 per group). Data are normalized to *Gapdh*. **(E)** Assessment of locomotor activity and exploratory behavior using the open field test (OFT). Representative tracking plots (up) and summary quantification of total distance traveled and number of center entry times (down) are shown for the different treatment groups (n = 10 per group). **(F)** Rotarod test quantification showing motor coordination and endurance in mice (n = 5 per group). Data in quantification plots are presented as mean ± SEM. Statistical significance was determined by one-way ANOVA and two-tailed unpaired t-test.

**Fig. 7.**
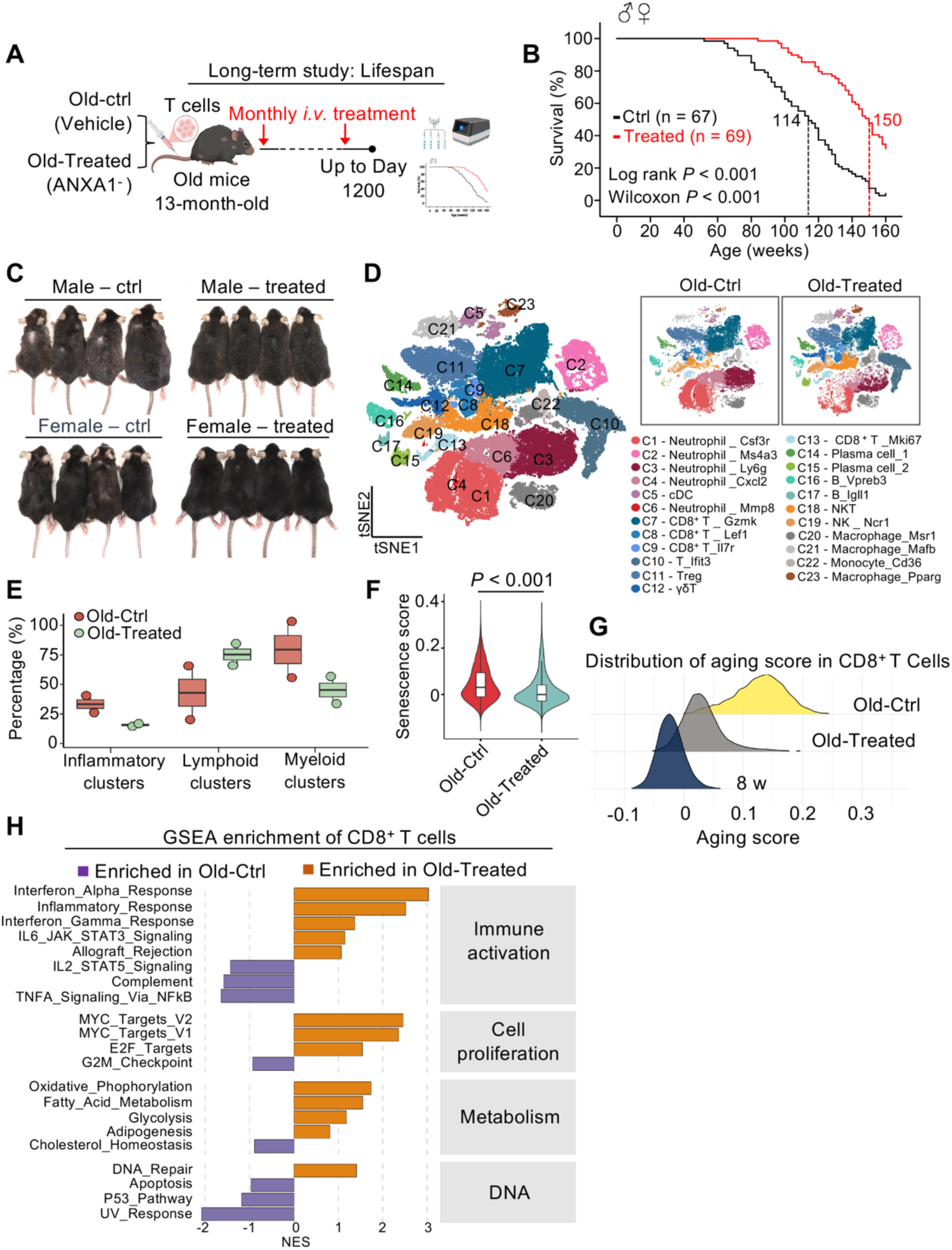
Long-term transfer of ANXA1⁻ CD8⁺ T cells ameliorates age-related phenotypes and reshapes the immune microenvironment in aged mice. **(A)** Schematic of the long-term therapeutic study design. Old (13-month-old) mice received monthly intravenous (*i.v.*) transfers of Anxa1⁻ CD8⁺ naïve T cells (Old-Treated group) or a vehicle control (Old-Ctrl group) for the duration of the study. **(B)** Kaplan-Meier survival curves for all mice with sexes combined. Statistical significance was determined by the Log-rank test. Median survival is indicated by dashed lines. **(C)** Representative photographs of mice at 30 months of age, highlighting the improved physical condition in the treated mouse. **(D)** Single-cell t-SNE projections of total CD45⁺ immune cells from the bone marrow of Old-Ctrl and Old-Treated mice (n = 2 per group). Left: Integrated UMAP colored by major cell lineages. Right: The same UMAP split by treatment group to visualize compositional shifts. **(E)** Box plots show the relative abundance of Inflammatory, Lymphoid, and Myeloid cell clusters as a percentage of total CD45⁺ cells between the two groups. **(F)** Violin plots comparing the senescence score, calculated per cell, between the Old-Ctrl and Old-Treated groups. The gene list for this score is provided in the Methods section. Statistical significance was determined by two-tailed unpaired t-test. **(G)** Density plot comparing the “Aging score” for all CD8⁺ T cells from the three experimental groups. The gene list for this score is provided in the Methods section. **(H)** GSEA enrichment analysis comparing Old-Treated versus Old-Ctrl CD8⁺ T cells. Bar plot shows NES for selected Hallmark gene sets. Purple bars indicate pathways enriched in Old-Ctrl, while orange bars indicate pathways enriched in Old-Treated cells.

The short-term study was designed to assess specificity, incorporating control groups receiving vehicle, ANXA1⁺ CD8⁺ naïve T cells, or unselected total CD4⁺/CD8⁺ T cells. Analysis revealed that only mice treated with ANXA1⁻ CD8⁺ naïve T cells—irrespective of whether the cells were sourced from young (Y-ANXA1⁻) or old (O-ANXA1⁻) donors—exhibited significant reductions in systemic senescence markers. This included lower p16^INK4a^ protein levels and decreased *Il6* mRNA expression in the spleen compared to aged controls (**Fig. 6B, D**). Furthermore, immunofluorescence analysis demonstrated reduction in p16^INK4a^-, and p53-positive senescent cells in the heart exclusively in mice treated with ANXA1⁻ CD8⁺ naïve T cells (**Fig. 6C**). These molecular improvements were accompanied by enhanced physical function. Recipients of ANXA1⁻ T cells showed enhanced exploratory behavior and movements in the open field test (**Fig. 6E**), along with improved motor coordination and endurance in the rotarod test (**Fig. 6F**). Notably, the therapeutic benefits observed from young and aged ANXA1⁻ T cells were comparable, and no significant improvements over vehicle controls were observed in mice receiving ANXA1⁺ T cells or total T cells (**Fig. 6B–F**). These findings highlight that the beneficial effects observed in this short-term context are specific to the ANXA1⁻ CD8⁺ naïve T cell population and independent of donor age.

We next investigated the long-term impact on healthspan and lifespan of ANXA1⁻ CD8⁺ naïve T cell transfer in naturally aged mice. A critical consideration for cell therapy is the persistence and phenotypic stability of transferred cells within the host environment. We established a long-term treatment regimen where aged (13-month-old) mice received monthly infusions of ANXA1⁻ CD8⁺ naïve T cells or vehicle control (**Fig. 7A**). Longitudinal tracking using CD45 congenic markers confirmed sustained engraftment of donor-derived cells in peripheral blood (**Fig. S9A**). Importantly, a substantial proportion of ANXA1⁻-derived cells retaining a naïve-like (CD62L⁺CD44⁻) phenotype and remaining largely surface ANXA1-negative at the protein level after 12 months transfer treatment (**Fig. S9B**). This sustained engraftment and phenotypic stability translated into significant therapeutic benefits. Kaplan-Meier analysis revealed a remarkable extension of lifespan in the treated cohort, with median survival increased by over 30 weeks compared to controls, observed consistently in both sexes (**Fig. 7B, Fig. S9A**). Beyond longevity, the ANXA1⁻ T cell therapy led to broad healthspan improvements. Treated mice maintained a visibly more youthful appearance at 30 months of age (**Fig. 7C**) and exhibited improvements across multiple physiological systems. These included improved skin structure (**Fig. S9C**), preserved trabecular bone mineral density (**Fig. S9D**), enhanced motor coordination and balance in the pole test (**Fig. S9E)**, and better cardiac function, as evidenced by increased left ventricular ejection fraction (LVEF) measured via echocardiography (**Fig. S9F**).

We further performed scRNA-seq profiling on bone marrow CD45⁺ immune cells from mice at Day 300 of the long-term therapeutic study. This analysis revealed that the overall immune cell composition shifted from the profile characteristic of aged controls towards to a state more reminiscent of younger animals, with lower frequency of myeloid lineage and inflammatory cluster (**Fig. 7D**). This systemic rejuvenation was evidenced by reduction in a calculated cellular ‘Senescence Score’ across the immune compartment (**Fig. 7F**). In-depth analysis of the host’s CD8⁺ T cells further demonstrated that the transcriptional state and subset distribution of host CD8⁺ T cells in treated mice diverged significantly from aged controls, partially recapitulating features observed in young mice (**Fig. S10A-E**). This rejuvenation was quantifiable as the decrease in a transcriptomic “Aging Score” and a concomitant increase in the “Stemness Score” of host CD8⁺ T cells (**Fig. 7G, Fig. S10F**). GSEA result further confirmed that treatment suppressed pro-inflammatory pathways and revived metabolic and proliferative gene programs in host CD8⁺ T cells (**Fig. 7H**). Collectively, these data indicate that long-term therapy with ANXA1⁻ CD8⁺ naïve T cells not only provides lifespan extension but also induces profound rejuvenating effects on the aged host itself.

## DISCUSSION

During the decades, pharmaceutical research has achieved notable advancements in developing targeted therapies for various aging-associated impairments, including cognitive defects and cardiovascular disorders(*24, 44–46*). A series of senolytic agents have been identified that can selectively eliminate senescent cells by targeting anti-apoptotic pathways that senescent cells rely upon(*44*). However, the therapeutic effects of these approaches to mitigate morbidity and mortality stemming from aging have been fairly restricted, further compounded by considerable side effects(*47, 48*). Partial inhibition of the metabolic checkpoint kinase and rapamycin target TOR has shown substantial promise, yet even low doses of rapamycin impair the generation of effector T cells in mice(*49*). This led to another exploration into finding the presence of pro-youthful circulating components intrinsic to the young host itself. Researches utilizing the HPB murine model have provided compelling evidence that exposure to a youthful systemic milieu can rejuvenate age-related impairments. Enhanced improvements were observed across tissues including muscle(*50*), liver(*28*), heart(*51*), and brain(*52*), while pathology of some age-related diseases such as Alzheimer’s disease also showed amelioration(*29, 53*). The precise molecular factors driving rejuvenation within the milieu of young blood have yet to be comprehensively identified.

In this study, we provide a comprehensive, multi-omic atlas of the aging human bone marrow, leading to the discovery of a novel, dysfunctional CD8⁺ naïve T cell subset defined by the surface expression of Annexin A1 (ANXA1). We establish ANXA1 as an intrinsic, functional brake on T cell cytotoxicity and demonstrate that the targeted adoptive transfer of ANXA1⁻ CD8⁺ naïve T cells acts as a potent senolytic therapy. Infusion of ANXA1^−^ CD8^+^ naïve T cells once a month into middle-aged mice significantly enhances multiple aspects of healthspan and reverses the progression of aging-related phenotypes, including those seen in locomotor function, cognition and cardiac health. The comprehensive therapeutic effects we observe are comparable to those observed in HPB mice models, confirmatory with aging phenotypes of multiple tissues, such as the muscle, heart, and brain, were ameliorated. Additionally, we demonstrate prolonged lifespan, with over 50% of treated mice surviving beyond 30 months of age, reaching similar even better extension effects(*46, 54*), equally 100 years in human lifespan, robustly indicative of the intermediary functions of immune factors in mediating youth milieu-induced rejuvenation of aged organisms. This fortifies the notion that reciprocal modulation exists between immune health and lifespan longevity, particularly regarding how the former can influence overall health. Herein, we present strong evidence complementing previous unidirectional research, partly substantiating the capacity of immunosenescence to disseminate senescence to peripheral organs(*2, 7*).

In this study, we move beyond the classical view of quantitative attrition and repertoire contraction to reveal a profound qualitative decline within the remaining naïve T cell pool(*55, 56*). The identification of the ANXA1⁺ subset as a distinct, phenotypically naïve population with a senescence-like, pro-inflammatory transcriptome represents a paradigm shift. It suggests that the naïve compartment is not a homogenous population of resting cells, but rather a landscape of functionally diverse subsets where aging progressively fosters the accumulation of a deleterious, “senescence-like” population. This finding converges with other recent human aging atlases that have identified robust, age-related transcriptional reprogramming in peripheral T cells and noted the emergence of ANXA1-expressing T cell subsets in the periphery and thymus (*17–19*). This ANXA1⁺ state, characterized by impaired proliferative potential, downregulation of T cell activation pathways, and an active SASP-like secretory program, positions these cells not as passive bystanders, but as active contributors to the cycle of inflammaging. Indeed, ANXA1’s role appears highly context-specific; for instance, while associated with dysfunction in our aging model, its overexpression has been reported to exert protective effects in other settings, such as attenuating autoimmune retinal inflammatory disease through Th17 cells(*57*) and suppressing the adipogenesis in aplastic anaemia(*58*). These disparate findings highlight the importance of investigating ANXA1 function specifically within the unique physiological context of naïve T cell aging, as explored in our study.

A central conclusion of our study is that ANXA1⁻ CD8⁺ naïve T cells function as a powerful form of endogenous senolytic therapy. There have been attempts using the adoptive transfer model to decipher age-related changes in immune cells, with contradictory results(*7*). A one-time infusion of young cells can be driven to exhaustion by an aged environment in mice, potentially due to unconventional interplay between the senescent milieu and adaptive immunity. Notably, in this study, we initiated interventions at the middle-aged stage of the murine lifespan and detected increased proportions of donor cells in the blood after multiple interventions, which elicited beneficial effects in aged mice. This discrepancy may arise from more severe comorbidities occurring later in life, while the sustained therapy can effectively delay age-related deterioration in late-life stages. The long-term adoptive transfer that we used in this study did not merely “replace” old cells with new ones; it actively remodeled the recipient’s bone marrow niche, shifting the landscape towards a more youthful state. The results that the host’s endogenous CD8⁺ T cells also exhibited a reduction in their “Aging Score” and an increase in their “Stemness Score” suggests that our therapy breaks a vicious cycle. By removing a source of pro-inflammatory SASP (the senescent cells), the ANXA1⁻ CD8⁺ naïve T cells likely create a less inflammatory, “pro-youth” niche that fosters the restoration of healthier hematopoiesis and function in the host’s own immune cells. This “bystander” rejuvenation effect has significant therapeutic implications, suggesting that periodic immunomodulation can trigger a durable, self-sustaining reversal of immunosenescence.

Together, our in-depth sequencing analyses on CD45^+^ cells from bone marrow have helped us elucidate the dynamic orchestration of the human immune compartment during aging. We identify ANXA1^+^ CD8^+^ naïve T cells as pivotal coordinators in aging, which possesses translational value for developing diagnostic assays to facilitate early clinical identification of individuals at risk for immune frailty or aberrant hyper-immunity. Evaluation of variation in the naïve T cell pool may also inform prognostic predictions and personalized therapeutic decisions to restore immune homeostasis.

Moreover, through cell-based therapy, we pioneer the concept of targeted immunomodulation as a novel anti-aging therapeutic approach, most importantly in extending healthspan. However, limitations in this study include insufficient discussion on TCR variations in CD8^+^ naïve T cells and the use of mice in specific pathogen-free (SPF) conditions. Going forward, whether and to what extent this T cell-based modulation benefited from enlarged TCR repertoire necessitates further investigation using different strains of mice.

## Acknowledgments

The authors would like to thank Y. Wang (Shanghai General Hospital) for help with cardiac functional assays; Z. Wei (Shanghai Sixth People’s Hospital) for help with micro-CT; L. Wang and K. Huang (Department of Laboratory, Shanghai General Hospital) for providing human PBMCs.

## Finding

National Natural Science Foundation of China Original Exploration Program (No. 82450903)

Shanghai Municipal Health Commission of Collaborative Innovation Cluster Program (No. 2024XJQ02)

National Natural Science Foundation of China (No. 82522076)

National Natural Science Foundation of China (No. 82373470)

National Natural Science Foundation of China (No. 82203914)

National Natural Science Foundation of China (No. 82504250)

Natural Science Foundation of Shanghai (No. 23ZR1480700)

## Author contributions

Conceptualization: H.W., Y.W., and S.G.; Investigation, Y.W., F. Z, F.L., Y.L, Y.S., Q. S., X.C., Z.W., X.Z., X.L., and M.G.; mouse strain construction and breeding, Y.L., Y.W., Q. Y., L. Z., and H.W.; data curation, Y.W., S.G, F.Z.; writing original draft, Y.W.; writing review and editing, H.W., Y.W., F.Z., F.L., and Y.L.; funding acquisition, H.W., F.L., Y.S., Z.W., X.Z., and S.D.; Supervision, H.W.; Project administration, H.W.

## Data and materials availability

All data needed to understand and assess the conclusions of this study are included in the Article and the Supplementary Information. scRNA-seq, scTCR-seq, and RNA-seq data supporting the findings of this study have been deposited in the Genome Sequence Archive (GSA) with accession number PRJCA023058.

## Supplementary Materials

### Materials and Methods

#### Human subjects

Human bone marrow samples were obtained during biopsy from healthy donors. Sample information included in scRNA-seq and scTCR-seq is provided in Supplementary Table 1. Human peripheral blood mononuclear cells (PBMCs) were obtained from blood buffy coats of healthy donors. Healthy, Asian, non-obese (BMI under 30) males and females were enrolled in the study. All donors provided written informed consent. This study was conducted in accordance with the principles of the Declaration of Helsinki and approved by the Research Ethics Boards of Shanghai General Hospital (Ethic review 2022-62, 2024KS75), Shanghai Sixth People’s Hospital (No. 2021-YS-093).

#### Animals

C57BL/6 CD45.2 mice, C57BL/6 CD45.1 mice, *Anxa1*^KO^ mice were purchased from Suzhou Cyagen Biotechnology Co., Ltd. affiliated to Cyagen US Inc. The mice were bred and maintained under specific pathogen-free (SPF) conditions. Mice were housed in cages with five mice per cage and kept on in a regular 12 h: 12 h-light: dark cycle. The temperature was 22 ± 1 °C and humidity was 40–70%. Age-matched and sex-matched mice were used for all of the experiments approved by the National Institutes of Health Guide for the Care and Use of Laboratory Animals with the approval (SYXK-2019-0028) of the Scientific Investigation Board of Shanghai General Hospital. To ameliorate any suffering that the mice observed throughout these experimental studies, the mice were euthanized by CO_2_ inhalation.

#### Cell culture

Mouse splenic CD8^+^ naïve T cells were isolated using MagniSort™ Mouse CD8 Naïve T cell Enrichment Kit (Invitrogen, cat. 8804-6825). For functional tests, 1×10^5^ cells were plated in RPMI 1640 medium (GIBCO cat. 11875093) supplemented with 10% fetal bovine serum (FBS, Gibco, cat: 10099141C) in the presence of plate-bound 2 μg/mL anti-CD3 antibody (Bio X Cell, clone 145-2C11) and 2.5 μg/mL anti-CD28 antibody (Bio X Cell, clone 37.51) and incubated for 12 h. The medium was refreshed every 2 days and the cells were sub-cultured according to the cell fusion. B16F10 cells (ATCC, Cat# CRL-6475) were cultured in high-glucose DMEM (Gibco, Cat# 11965092) supplemented with 10% fetal bovine serum (FBS; Gibco, Cat# 10099141C) and 1% Pen-strep (Gibco, Cat# 15240062) in a 37°C, 5% CO₂ incubator. Upon reaching 80% confluence, cells were washed with PBS and detached using 0.25% trypsin (37°C, 5% CO₂) for 1 min before subculturing or experimental use. Isolation and extraction of skin fibroblasts: Cutting the skin tissue blocks into 0.2 cm × 0.2 cm pieces. Place the pieces evenly in a six-well plate. Add 1 mL of high-glucose DMEM medium containing 20% FBS (Gibco, Cat# 10099141C) and 1% P/S to each well, ensuring the medium just covers the tissue pieces. Incubate the plate in a 37°C, 5% CO₂ incubator. Change the medium every other day. After two weeks, digest the cells and expand the culture to passage 3 (P3) using serum-free medium, then −80°C cryopreserve the cells.

#### Single-cell RNA sequencing

Fresh human and mouse bone marrow samples were placed in saline at 4 °C until further processing. Single-cell suspensions were generated by dissecting tissues and subjected to fluorescence-activated cell sorting (FACS) to exclude doublets, debris, and DAPI^+^ dead cells. CD45^+^ live cells were sorted, centrifuged and resuspended in PBS containing 0.04% BSA. Chromium Single Cell 3′ v3 (10x Genomics) library preparation was conducted by according to the manufacturer’s instructions in Shanghai Xurangene Biotechnology Co., Ltd. The resulting libraries were sequenced with an Illumina HiSeq 4000 platform. Trimmed data were processed using the CellRanger (10x Genomics, version 3.0) and further filtered, processed, and analyzed using the Seurat package(*59*) (version 4.3.0) with R (version 4.2.3). The Seurat R package was used for subsequent analysis. Cells with fewer than 200 genes, more than 5000 genes, or more than 5% mitochondria content were removed. Cells identified as doublets using DoubletFinder(*60*) R package were filtered out. The filtered data were normalized using a scaling factor of 10,000 to generate transcripts per kilobase million (TPM)-like values. Filtered samples were integrated using the FindIntegrationAnchors and IntegrateData functions with default parameters (dimensionality = 30). Most variable genes were selected using the FindVariableFeatures function and the genes were then used for principal component analysis (PCA). Clustering was performed using the FindClusters function with a resolution selected for different datasets. Results were visualized using the Seurat package and ggplot2 package. For human scRNA-seq analysis, we computationally filtered a sub-cluster corresponding to hematopoietic progenitor cells (*ITGA2B*, *KIT*) after initial clustering of human bone marrow sequencing, in order to focus our analysis on mature CD45⁺ immune cells. Resulting unbiased clustering of the cells was not affected by batch effects derived from ambient RNA or depth and quality of sequencing. The R package ggplot2, ComplexHeatmap, pheatmap were used to plot the results of differential expression analyses. Z-score normalization was performed to standardize gene expression. CD8^+^ T cells trajectory analysis was performed by Monocle. GO and GSEA were performed using R package clusterProfiler and MSigDB Hallmark gene sets. Mouse CD8^+^ T cells pre-clustered and renamed according to the marker genes were used as an input for Monocle 2 in the pseudo-time trajectory analysis(*61*). Genes with expression levels lower than 0.1 and genes expressed by fewer than 10 cells were removed. For quantifing the activity of specific biological pathways at the single-cell level, we used the AddModuleScore function from the Seurat R package (v4.3.0) to calculate module scores. Senescence Score was used to assess the overall senescence and Senescence-Associated Secretory Phenotype (SASP) signature of individual immune cells. The score was based on a set of canonical senescence and SASP-related genes derived from MSigDB (R-MMU-2559582). Aging Score (T cell aging score) was specifically used to assess the aging-related transcriptional signature of host CD8⁺ T cells. The gene set was defined based on genes upregulated in the Old-Ctrl vs. 8w DEG analysis from our dataset. Stemness Score was used to quantify the stemness or ‘youthful’ state signature of host CD8⁺ T cells. The gene set was defined based on canonical T cell stemness/memory-precursor markers(*62, 63*) including ‘*Tcf7*’, ‘*Sell*’, ‘*Lef1*’, ‘*Ccr7*’, ‘*Il7r*’.

#### Single-cell T Cell Receptor sequencing

Single-cell 5′ and V(D)J libraries were prepared following the protocol provided by the 10× genomics. In short, T cell suspensions were loaded on a Chromium Single Cell Controller (10× Genomics) to generate single-cell gel beads in emulsion (GEMs) using Chromium Single Cell V(D)J Reagent Kits. Captured cells were lysed, and the released RNAs were barcoded through reverse transcription in individual GEMs. Each single-cell 5’ and V(D)J libraries were sequenced by the Illumina Novaseq 6000 using 150 paired-end reads. Single-cell V(D)J data was processed using Cell Ranger (10× Genomics, version 3.1.0). Paired α and β CDR3 sequences from human bone marrow were pooled together to identify common clonotypes across samples. Data clustering and integrating were performed by scRepertoire(*64*) R package. To determine hyperexpanded TCR sequences identified in our scTCR-seq, the CDR3 region of each α and β chain was compared to the CDR3 repertoire from the VDJdb at https://vdjdb.cdr3.net/.

#### RNA-seq

For mouse splenic CD8^+^ naïve T cells and human PBMCs, samples were washed twice with PBS, and total RNA was extracted with RNAiso Plus (Takara Bio cat. 9109) and purified with magnetic oligo (dT) beads after denaturation. Purified mRNA samples were reverse-transcribed into fragmented DNA samples and adenylated at the 3′ ends. Adaptors were ligated to construct a library. DNA was quantified by Qubit (Invitrogen). DNA samples were sequenced by an Illumina HiSeq X Ten SBS instrument from Xurangene Bio (Shanghai). Raw data were converted into FASTQ format, and transcript per million fragments mapped (fragments per kilobase) was calculated and log2-transformed with Cuffnorm. Differential gene transcripts were analyzed with DESeq and enriched for the GO/Kyoto Encyclopedia of Genes and Genomes pathway. Pathway enrichment analysis was performed using the clusterProfiler(*65*) package.

#### Real-time quantitative PCR

Total RNA was isolated using RNAiso Plus (Takara Bio cat. 9109), quantified with a NanoDrop spectrophotometer, and reverse-transcribed into cDNA using an M-MLV First-Strand Synthesis Kit (Invitrogen, cat: C28025-021) with oligo(dT) primers. qPCR was conducted with the Hieff qPCR SYBR Green Master Mix (Yeasen Bio-tech, cat: 11202ES03) in a ViiA 7 Real-Time PCR system (Applied Bio- systems). The relative expression of target genes was confirmed using the quantity of target gene/quantity of GAPDH. The following primers were used:

*Gapdh* forward: 5’-TGTGTCCGTCGTGGATCTGA-3’,
*Gapdh* reverse: 5’-CCTGCTTCACCACCTTCTTGA-3’;
*Il6* forward: 5’-CCAAGAGGTGAGTGCTTCCC-3’;
*Il6* reverse: 5’-CTGTTGTTCAGACTCTCTCCCT-3’.

#### Western blotting

For immunoblotting, cultured cells were lysed in Cell lysis buffer for Western and IP (Beyotime cat. P0013) containing a protease and phosphatase inhibitor cocktail (Thermo Fisher Scientific cat. 78440). anti-CDKN2A/p16^INK4a^ antibody (abcam, cat. ab189034, 1:1000 dilution), GAPDH Mouse Monoclonal Antibody (Proteintech, cat. 60004-1-Ig-100ul, 1:10000 dilution), Alpha Tubulin Monoclonal antibody (Proteintech, cat. 66031-1-Ig, 1:1000 dilution), and HRP-labeled goat anti-mouse IgG (H + L) (Beyotime cat. A0216, 1:2000 dilution) were used as antibodies. The signal was detected with ECL Western Blotting Substrate (Thermo Fisher Scientific cat. 34095) and an Amersham Imager 600 (GE Healthcare). The images were cropped for presentation.

#### Isolation of human PBMCs

Venous blood samples were collected in Sodium-Heparin vacutainers. Peripheral blood mononuclear cells (PBMCs) were isolated using Lymphoprep™ (STEMCELL Technologies cat. 07811) according to the manufacture’s protocol. PBMCs were then washed twice in DPBS for further tests. In flowcytometric analysis on ANXA1 expression, cells were then washed and stained with Annexin V Binding Buffer (BioLegend cat. 422201) according to the manufacture.

#### Flow cytometry

Single-cell suspensions were pelleted and resuspended in PBS with 2% FBS containing fluorophore-conjugated antibodies. Cells were initially stained with antibodies targeting cell surface proteins and with Live/ dead Fixable Violet Dead Cell Stain Kit (Thermo Scientific, cat. L34964) for 30 min on ice and washed with PBS containing 2% FBS. For intra- cellular target staining, cells were then fixed and permeabilized using a Cytofix/Cytoperm kit (BD Biosciences, cat. 554714). For intracellular cytokine staining, single-cell suspensions were plated at 4 × 10^6^ cells per well in a 96-well round bottom plate, resuspended in 1× cell stimulation cocktail (plus protein transport inhibitors, eBioscience, cat. 00-4975-03) and incubated at 37 °C for 4 h before antibody staining. Samples were run on a BD Fortessa and analyzed using FlowJo software (version 10.6.2). The following antibodies and reagents were used for human and mouse flow cytometry: APC anti-human Annexin A1[Clone: 74/3] (BioLegend, cat: 831604), PE anti-human RANTES (CCL5) [Clone: VL1] (BioLegend, cat: 515503), PerCP/Cy5.5 anti-human CD45RA [Clone: HI100] (BioLegend, cat: 304122), Brilliant Violet® 605 anti-human CD62L [Clone: DREG-56] (BioLegend, cat: 304834), PE (Phycoerythrin)/Cy7® anti-human CD25 [Clone: BC96] (BioLegend, cat: 302611), APC anti-human CD45 [clone: HI30] (BioLegend, cat: 304012), PE anti-human CD3 [Clone: UCHT1] (BioLegend, cat: 300441), FITC anti-human CD8a [Clone: RPA-T8] (BioLegend, cat: 301050), anti-CDKN2A/p16^INK4a^ antibody (abcam, cat. ab189034), DAPI (BD Pharmingen, cat: 564907), PE (Phycoerythrin)/Cy7® anti-mouse CD69 [clone: H1.2F3] (BioLegend, cat: 104512), PE anti-mouse CD25 [Clone: PC61] (BioLegend, cat: 102008), Brilliant Violet 421 anti-mouse CD25 [Clone: PC61] (BD Pharmingen, cat: 562606), Alexa Fluor® 700 anti-mouse CD45.2 [Clone: 104] (BioLegend, cat: 109822), PE anti-mouse CD45.1 [Clone: A20] (BioLegend, cat: 110708), PE-Cy7 anti-mouse IFN-γ [Clone: XMG1.2] (BD Pharmingen, cat: 557649), APC/Cyanine7 anti-mouse CD279 (PD-1) [Clone: 29F.1A12] (BioLegend, cat: 135223), Brilliant Violet 650 anti-mouse CD223 (LAG-3) [Clone: C9B7W] (BioLegend, cat: 125227), Annexin A1 Monoclonal antibody [Clone: 1E1B7] (Proteintech, cat: 66344-1-IG), Alexa Fluor® 488 Anti-Annexin A1/ANXA1 antibody [Clone: EPR19342] (abcam, cat: ab225513), PE anti-mouse/human CD44 [Clone: IM7] (BioLegend, cat: 103024), APC anti-mouse CD62L [Clone: MEL-14] (BioLegend, cat: 104412), PE Anti-Mouse CD107a/LAMP-1 Antibody [Clone: 1D4B] (Elabscience, cat: E-AB-F1254D), PE-CF594 Hamster Anti-Mouse CD3e [Clone: 145-2C11] (BD Pharmingen, cat: 562286), PerCP/Cyanine5.5 anti-mouse CD8a [Clone: 53-6.7] (BioLegend, cat: 100734). For proliferation assay, CellTrace™ Violet (Invitrogen, cat: C34557) was used according to the manufacture’s protocol.

#### Anti-tumor assays

B16-F10 melanoma cells (ATCC, CRL-6475) were cultured in high-glucose DMEM (Gibco, cat. 11965092) with 10% FBS (Gibco, cat. 10099141C) and 1% Pen-strep (Gibco, cat. 15240062). C57BL/6J mice (8–10 weeks old) were subcutaneously (s.c.) injected on the right flank with 2 × 10⁵ B16-F10 cells in 100 µL PBS. For T cell preparation, CD8⁺ T cells were isolated from spleens of WT and *Anxa1*KO mice using MagniSort™ Mouse CD8 T cell Enrichment Kit (Invitrogen, cat. 8804-6822-74). On day 7 post-tumor inoculation (when tumors were palpable), mice received 2 × 10⁶ CD8⁺ T cells (WT or *Anxa1*KO) via intravenous (*i.v.*) injection. Tumor volume was measured with calipers every 2–3 days (Volume = [Length × Width²]/2). Mice were euthanized 14 days post-transfer, and tumors were excised and weighed.

For invasion assay, U-2 OS osteosarcoma cells (ATCC, HTB-96) were cultured in McCoy’s 5A medium (10% FBS). T cell-mediated inhibition of invasion was assessed using 24-well (8.0 µm pore) Transwell inserts (Corning) coated with Matrigel (1:8 dilution). Surface Anxa1⁻ and Anxa1⁺ CD8⁺ naïve T cells were sorted from young (8-week-old) and aged (13-month-old) mice using MagniSort™ Mouse CD8 Naïve T cell Enrichment Kit (Invitrogen, cat. 8804-6825). The Transwell lower chamber contained 600 µL RPMI (Hyclone, cat. SH30027.FS) with 10% FBS as a chemoattractant. U2OS cells (5 × 10⁴) were co-seeded with T cells (2.5 × 10⁵; E:T ratio 5:1) in 200 µL serum-free medium into the upper chamber. Control wells contained only U2OS cells. After 24h incubation, non-invading cells on the upper surface were removed. Invading cells on the lower surface were fixed (4% PFA, YOBIBIO, cat. YB1001), stained (0.1% crystal violet), and imaged. Invasion was quantified by counting cells from at least five random fields per membrane.

#### Senescence induction

Mouse primary fibroblasts were seeded 48 h before UV-B irradiation. The culture medium was removed, and PBS was added to the cells before UV-B irradiation (200 mJ m^−2^) as described previously(*66*). Following irradiation, the prewarmed culture medium containing the respective treatment cells was added to the flask. Cells were cultured for 5 days to develop a senescent phenotype before further analysis. Doxo was used to induce senescence in mouse by intraperitoneal injection (20mg/kg). Etoposide (MCE, HY-13629) was used at 10 μM for 24 h to mouse primary fibroblasts. Lentivirus particles were obtained by co-transfecting PGMLV-CMV-MCS-3×Flag-EF1-ZsGreen1-T2A-Puro overexpression plasmid with packaging plasmids (pMD2.G and psPAX2) into HEK293T cells. Human primary fibroblasts were infected with the above lentivirus particles and then were sorted for GFP^+^ cells to use for subsequent experiments.

#### Adoptive transfer of mouse model

CD45.1^+^ CD8^+^ naïve T cells first were isolated from the spleen of young (4-6 weeks), or old (13-month-old) C57BL/6J mice using MagniSort™ Mouse CD8 Naïve T cell Enrichment Kit (Invitrogen, cat. 8804-6825) and then ANXA1^−^, ANXA1^+^ live cells were sorted within for further transfer. CD45.1^+^ CD4^+^ T cells were isolated from the spleen of young (4-6 weeks) C57BL/6J mice using MagniSort™ Mouse CD4 T cell Enrichment Kit (Invitrogen, cat. 8804-6821-74), CD45.1^+^ CD8^+^ T cells were isolated from the spleen of young (4-6 weeks) C57BL/6J mice using MagniSort™ Mouse CD8 T cell Enrichment Kit (Invitrogen, cat. 8804-6822-74). 2 × 10^6^ cells donor mice were *i.v.* injected into 13-month-old CD45.2^+^ recipient mice. Each month post cell transfer, CD45.1^+^ and CD45.2^+^ cells were monitored by flow cytometry in the blood.

#### Three-dimensional microcomputed tomography

X-ray computed micro-CT was performed using a SkyScan 1276 (Bruker micro CT, Belgium) equipment(*67*). In vivo (whole body and body fat) scanning parameters were: image pixel size of 35.85 μm, 60 kV, 300 μA, 0.5-mm aluminum filter, 2 frame averaging. For *in vitro* bone microarchitecture analysis, tibias were scanned using the following parameters: image pixel size of 8.96 μm, 50 kV, 450 μA, 0.5-mm aluminum filter. The 1.6 version of NRecon software (Bruker) was used for three-dimensional (3D) reconstruction, 3.3.0 version of CTvox software (Bruker)was used for 3D photos. The 1.13 version of CTan software (Bruker) was used for bone analysis.

#### Immunohistochemistry and immunofluorescence

Skin specimens were embedded in paraffin and sectioned at the Histology Core of the Shanghai General Hospital. Masson’s trichrome (Poly Scientific R&D; HT15-1KT, Sigma-Aldrich) staining was performed according to the manufacturer’s protocol. Immunohistochemistry slides were blindly quantified using ImageJ software. For immunofluorescence staining, OCT treated mouse tissues slices were fixed and blocked with blocking buffer (5% normal goat serum, 3% BSA, and 0.2% Triton X-100 in PBS) for 1 h and stained with primary antibody overnight at 4 °C. Afterward, the slices were rinsed three times in PBS and stained with relative secondary antibodies for 1 hour at RT. Nuclei were stained with DAPI (BD Biosciences, cat: 564907) at RT for 10 min. After being washed in PBS, the slices were mounted (Beyotime Biotechnology, cat: P0126) and visualized under a confocal microscopy system (Leica, TSC SP8). For immunofluorescence staining of cells, suspending cells were resuspended in PBS and centrifuged onto slides at 1,000 rpm for 5 min, while adherent cells were grown directly on coverslips. Cells were washed 3 times with PBS, fixed with 4% PFA for 30 min, and permeabilized with 0.4% Triton X-100 for 20 min. After PBS washes, cells were blocked with 5% goat serum for 1 h at room temperature followed by primary antibody against p16^INK4a^, incubation overnight at 4°C. The next day, samples were incubated with fluorochrome-conjugated secondary antibodies (all from Life Technologies, 1:1,000 dilution) for 1 h at room temperature. Primary antibodies include: anti-CDKN2A/p16^INK4a^ antibody (abcam, cat. ab189034), anti-p21 antibody [EPR18021] (abcam, cat. ab188224), p53 (1C12) Mouse mAb (Cell Signaling, cat. 2524S), anti-γH2AX (phospho S139) (abcam, cat. ab81299), Alexa Fluor® 488 Anti-Annexin A1/ANXA1 antibody [EPR19342] (abcam, cat: ab225513), Granzyme B Polyclonal antibody (Proteintech, cat. 13588-1-AP).

#### Open field test

Locomotor activity and exploratory cognition were tested using open field test as previously reported(*68*). In short, an open field apparatus was used, and the central zone was a square 20 cm away from the wall of the apparatus. Mice were placed in the central zone and allowed autonomous exploration. Before formal tests, mice were habitude to the apparatus for 2 days to eliminate potential anxiety as previous described. The time of movement and spent in the central zone along with whole trajectory were recorded for 5 min each time. The apparatus was cleaned after each test.

#### Rotarod test

Motor coordination was assessed employing an automated rotating rod apparatus following established protocols(*69*). Individual subjects were positioned in designated compartments of the rotating cylinder, which accelerated uniformly at 8 rpm/min from an initial velocity of 4 rpm until reaching 44 rpm. Trials terminated after three consecutive falls per subject during training. The formal test was conducted on the fourth day, with each subject undergoing a single trial to record the duration of maintaining balance on the rotating rod.

#### Pole test

Pole tests were performed in testing. A 75 cm long with a 9 mm diameter-sized metal rod wrapped with bandage gauze was used. Mice were trained in three test trials for 2 days. The time to turn and total time to place all four paws on the base were measured after placing the mice from the top of the pole. The pole was cleaned after each trial.

#### Cardiac functional characterization

Briefly, male and female mice were anesthetized with 1% isoflurane at a rate of 1.0 L/min. MyLab 30CV machine (Esaote SpA) with a 10-MHz probe was used to obtain two-dimensional echocardiograms. Mice were placed supine on the platform, and the transducer was used to obtain Parasternal Long-Axis View (PLAX).

#### Quantification and statistical analysis

The data were analyzed using GraphPad Prism (version 9) and are presented as the mean ± SEM. A Student’s *t*-test was used to compare two conditions, and an analysis of variance (ANOVA) with Bonferroni or Newman–Keuls correction was used for multiple comparisons. A simple linear regression model was used to analyze the correlations using R software. The probability values of <0.05 were considered statistically significant. The error bars depict the SEM. Illustrative images were created in https://BioRender.com.

**Fig. S1.**
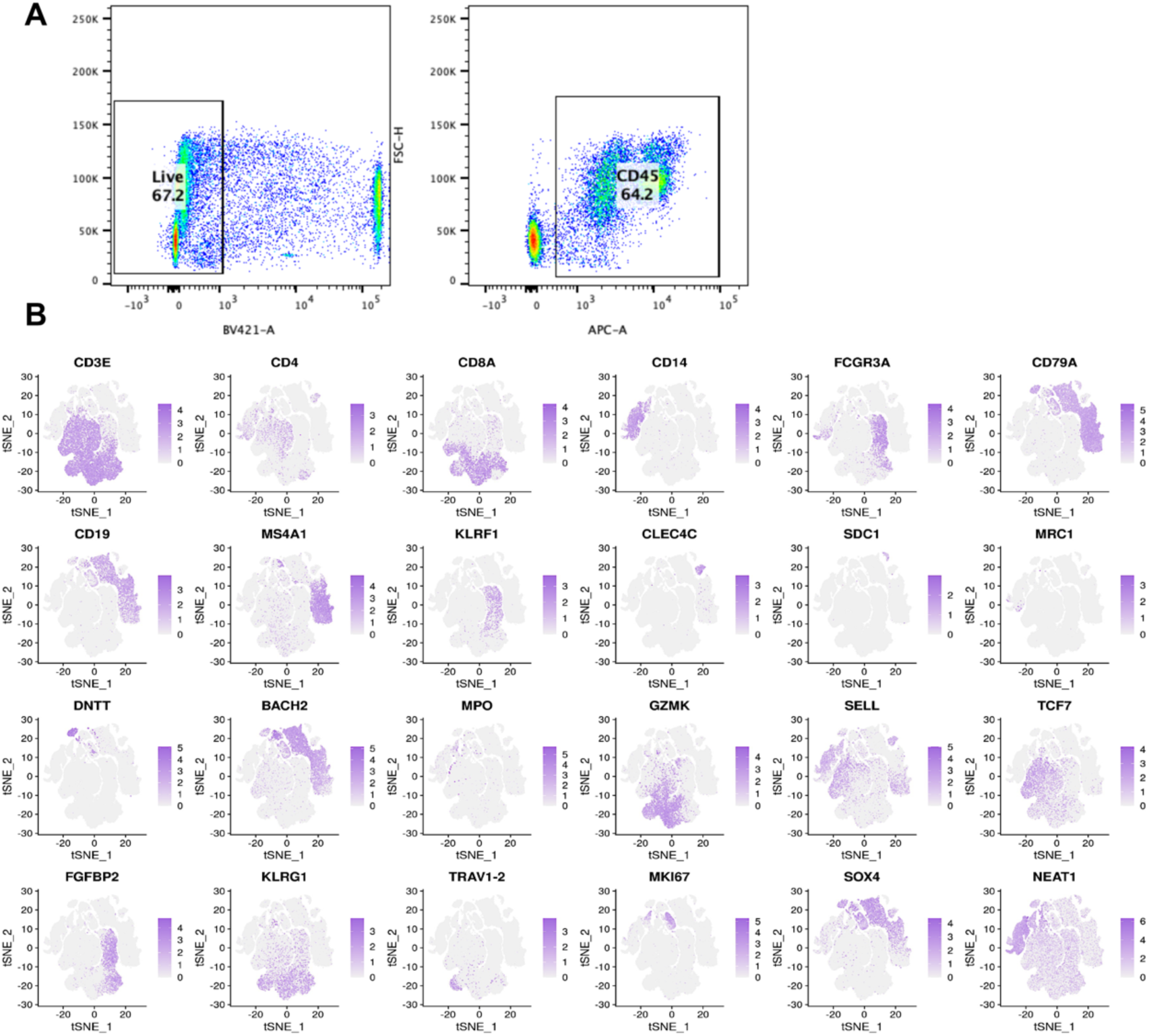
Quality control and cell lineage annotation of human bone marrow scRNA-seq data. **(A)** FACS gating strategy for isolating CD45⁺ immune cells from human bone marrow for scRNA-seq and scTCR-seq analysis. **(B)** Feature plots of canonical marker genes used to identify the major immune cell populations annotated in Fig. 1B.

**Fig. S2.**
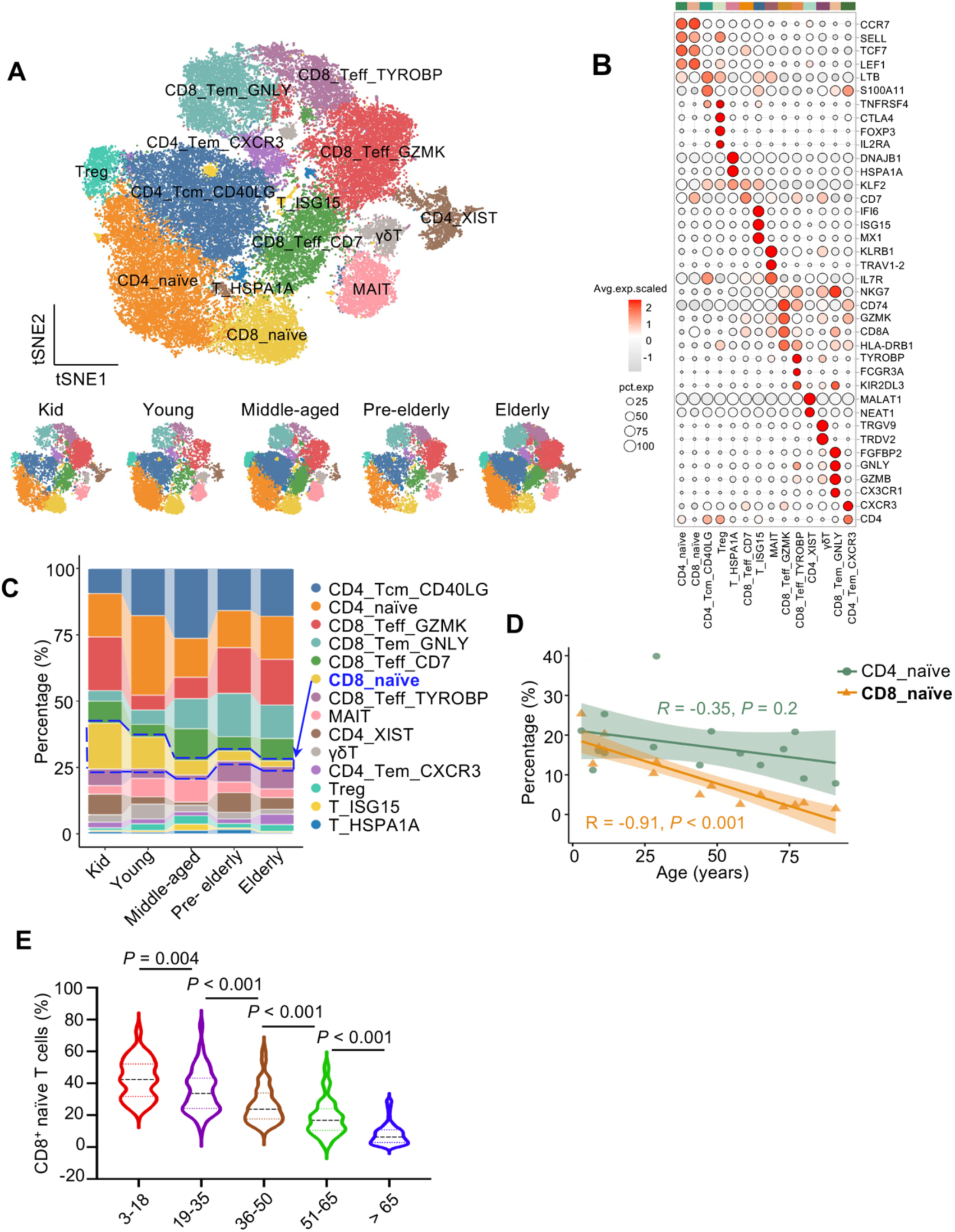
scRNA-seq analysis of T cells in human bone marrow samples. **(A)** T-SNE clustering of all identified T-cell subsets from human bone marrow for scRNA-seq data. T cells from different groups of donors are shown separately. **(B)** Dot plot on signature genes identified in each T cell subsets. Color scale represents the average expression level of selected genes and dot size represents the proportion of cells expressing marker genes in each cluster. Names of each cell type are listed below, corresponding with those in (**A**). **(C)** Stacked bar plot showing the proportional changes of cell clusters between 5 age groups. **(D)** Correlation between the proportion of naïve T-cell subsets and donor age. Scatter plots show the percentage of CD4⁺ and CD8⁺ naïve T cells relative to total T cells versus donor age. Solid lines indicate linear regression fits, with Pearson correlation coefficient (R) and *P*-values displayed. **(E)** Validation of age-related decline in CD8⁺ naïve T-cell frequency in PBMCs. Box plot showing the percentage of CD8⁺ naïve T cells (identified by flow cytometry as in Fig. 2E) within total CD8⁺ T cells across age groups in the large validation cohort (n=388, same cohort as Fig. 2F).

**Fig. S3.**
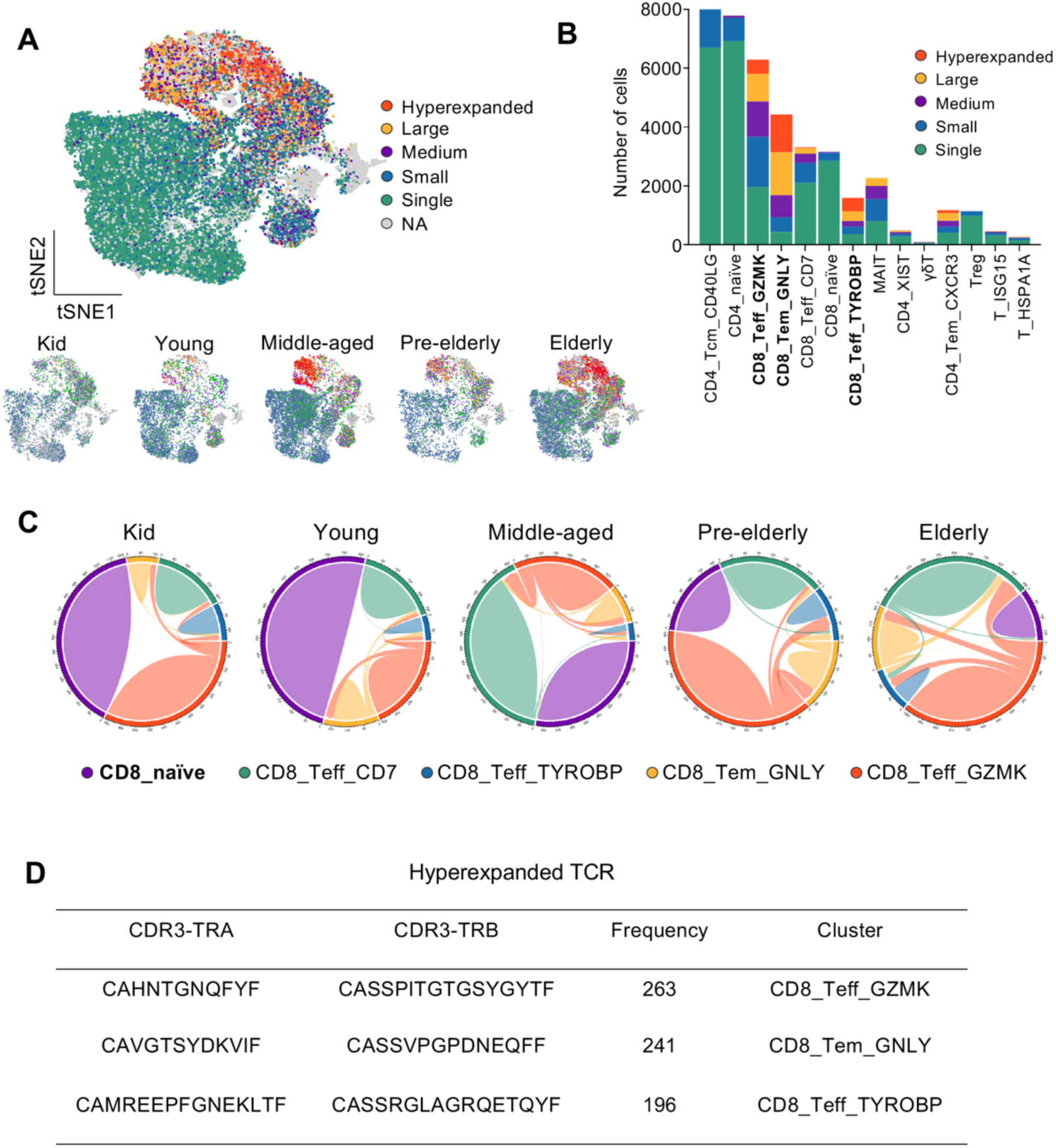
scTCR-seq analysis of the T-cell receptor repertoire. **(A)** T cell receptor (TCR) clone-types analysis in scTCR-seq (n=13) projected on t-SNE plot in Fig. S2A. Colors represent different clonotypes based on numbers (x) of same clonotype. 6 levels are defined, including: hyperexpanded (100 < x ≤ 500), large (20 < x ≤ 100), medium (5 < x ≤ 20), small (1 < x ≤ 5), single (x = 1) and NA. Separated clustering plots are shown below from samples of different age groups. **(B)** Distribution of 5 clonotypes in all cell clusters. Bold texts represent clusters with different levels of clonotypes, especially with larger expanded clones **(C)** Chord diagram shows the repertoire changes in CD8^+^ T cells compared between samples in different age groups. **(D)** Top 3 hyperexpanded TCR-CDR3 sequences are listed, all enriched in CD8^+^ T cells.

**Fig. S4.**
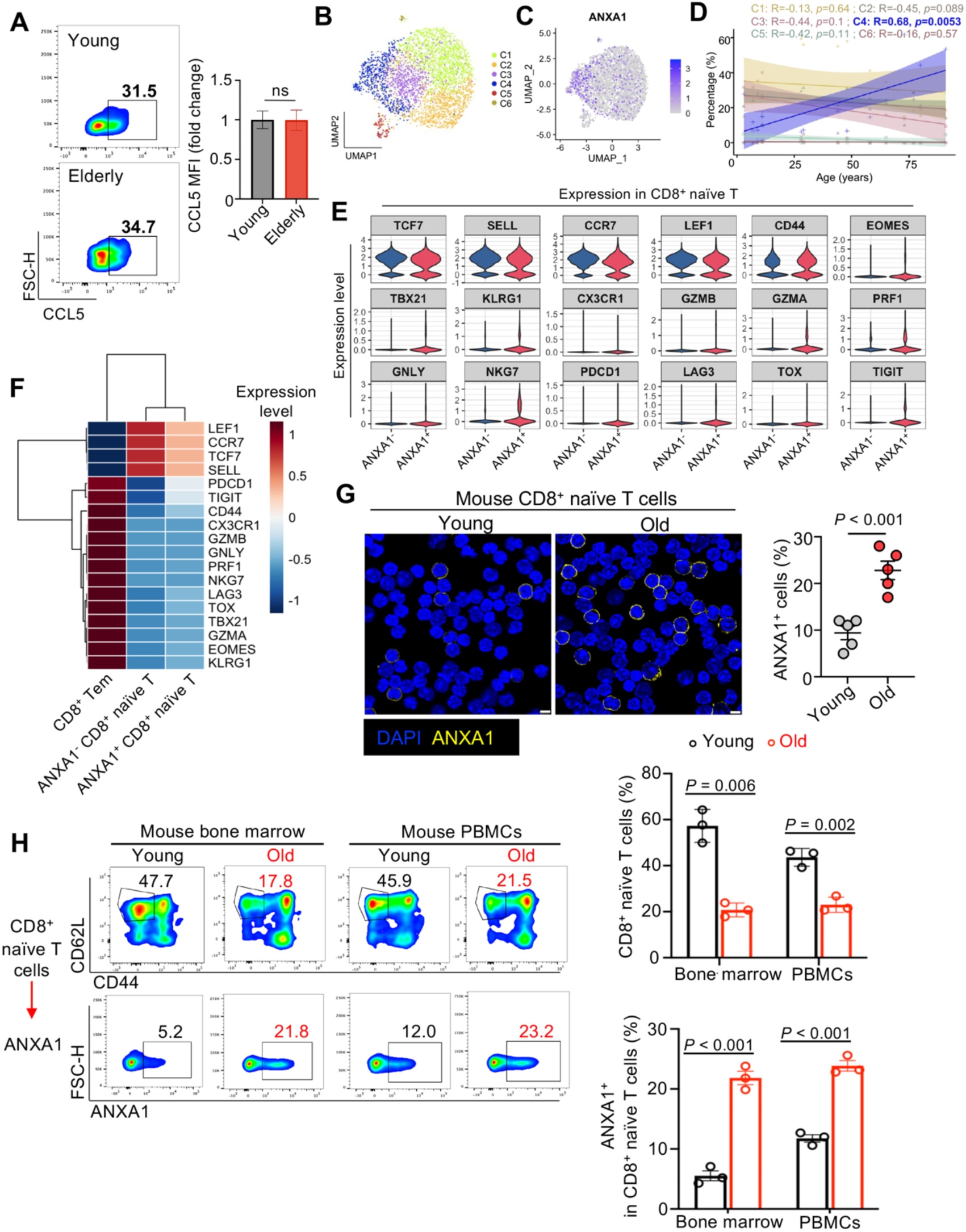
Phenotypic definition and validations of ANXA1^−^ and ANXA1⁺ CD8⁺ T cells. **(A)** Representative flow cytometric analysis of CCL5 expression on CD8^+^ naïve T cells in samples of young (19-35 y) and old (>65 y) group (n= 5 per group). **(B)** Sub-clustering of CD8⁺ naïve T cells from the scRNA-seq dataset. UMAP projection showing distinct subclusters (C1-C6) identified within the CD8⁺ naïve T cell population. **(C)** Feature plot overlaying normalized ANXA1 expression onto the UMAP projection from (**B**), highlighting its specific enrichment in the C4 subcluster. **(D)** Scatter plots showing the correlation between the relative abundance of each subcluster (as a percentage of total CD8⁺ naïve T cells) and donor age. Lines represent linear regression fits; Pearson correlation coefficient (R) and *P*-values are indicated. (**E** and **F**) Violin plots (**E**) and heatmap (**F**) validating the naïve identity of both ANXA1^−^ and ANXA1⁺ CD8⁺ naïve T cells defined in human scRNA-seq data. Average expression of canonical lineage markers for naïve, memory, effector, and exhausted T cells is compared across three populations with CD8⁺ effector memory (Tem) cells as a non-naïve control. Expression values are scaled per row. **(G)** Representative immunofluorescence images and quantification of ANXA1 surface expression on mouse CD8⁺ naïve T cells from Young (8-week-old, n = 5) and Old (13-month-old, n = 5) mice. Scale bar, 5 µm. **(H)** Representative flow cytometry results and quantifications on the percentage of CD8^+^ naïve T cells and surface ANXA1 expression in young and old mice (n = 3 per group). Data in quantification plots are presented as mean ± SEM. Statistical significance was determined by one-way ANOVA and two-tailed unpaired t-test.

**Fig. S5.**
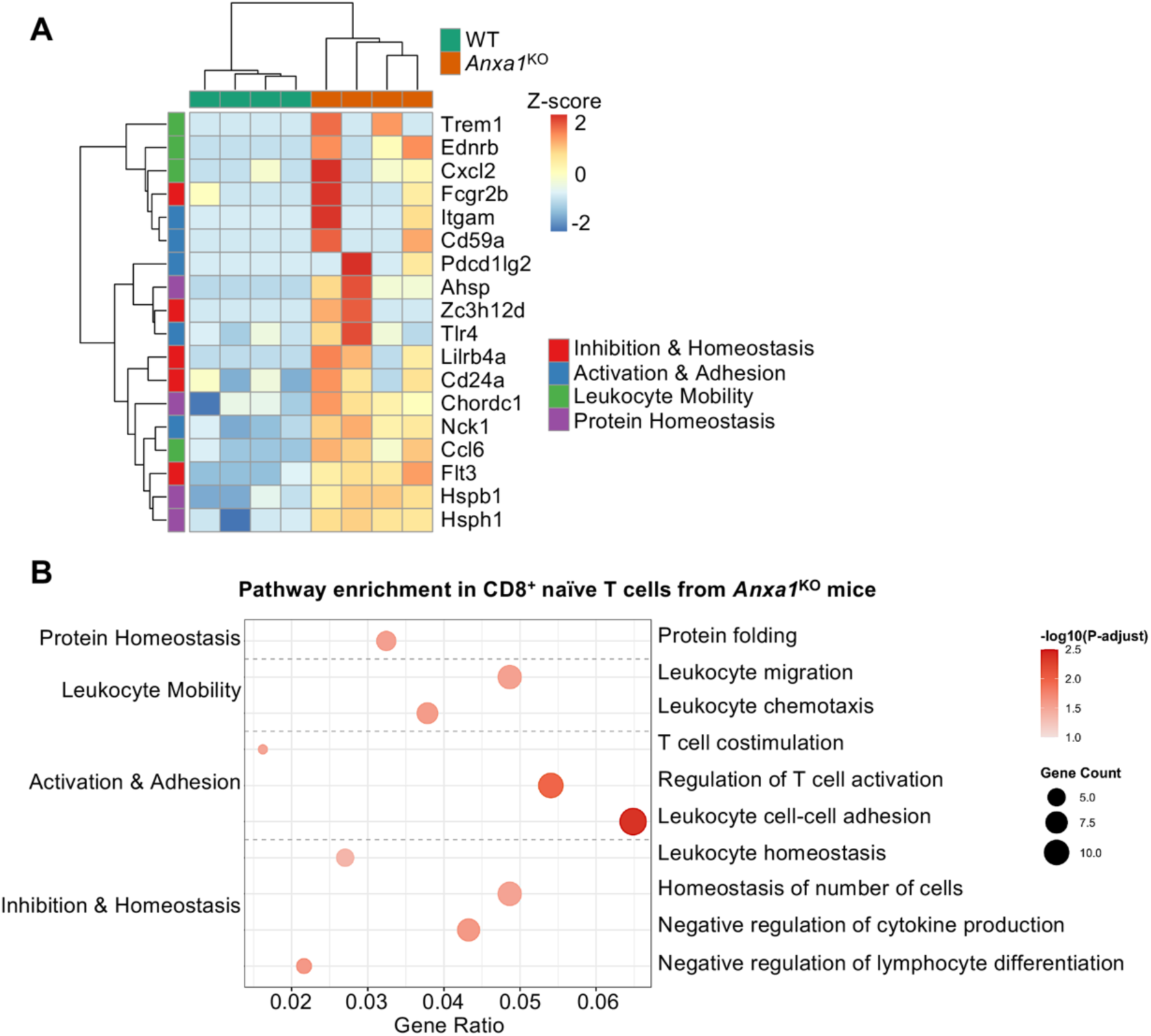
ANXA1-deficient CD8⁺ naïve T cells exhibit a primed phenotype at basal state. **(A)** Heatmap of DEGs from RNA-seq of resting WT and *Anxa1*^KO^ CD8⁺ naïve T cells. **(B)** GO enrichment bubble plot of pathways upregulated in *Anxa1*^KO^ versus WT CD8⁺ naïve T cells at basal state.

**Fig. S6.**
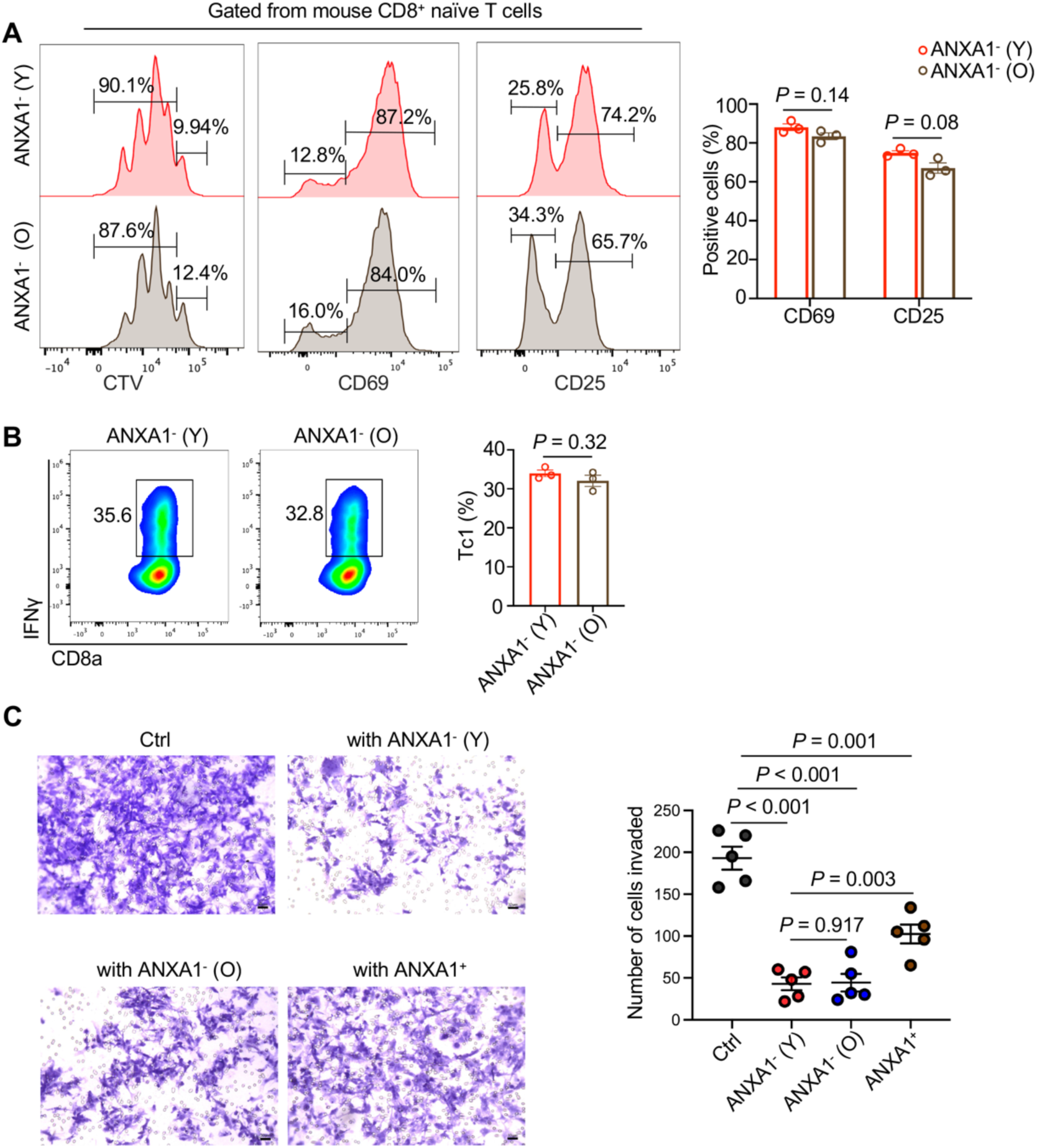
ANXA1^−^ CD8^+^ naïve T cells cells from young and old mice are functionally comparable *in vitro*. **(A)** Representative flow cytometry histograms and quantifications of CTV-staining, CD69 and CD25 expression of ANXA1^−^ CD8^+^ naïve T cells from young and aged mice (n = 3 per group). **(B)** Representative flow cytometry results and quantifications of Tc1 differentiation of ANXA1^−^ CD8^+^ naïve T cells from young and aged mice. **(C)** Representative images and quantification from a transwell invasion assay showing that both young and old mouse ANXA1⁻ T cells, but not ANXA1⁺ T cells (n = 5 per group), significantly inhibit the invasion of U-2 OS tumor cells, with arrows indicate tumor cells passed the transwell membrane. Data are presented as mean ± SEM. *P*-values were determined by one-way ANOVA and two-tailed unpaired t-test.

**Fig. S7.**
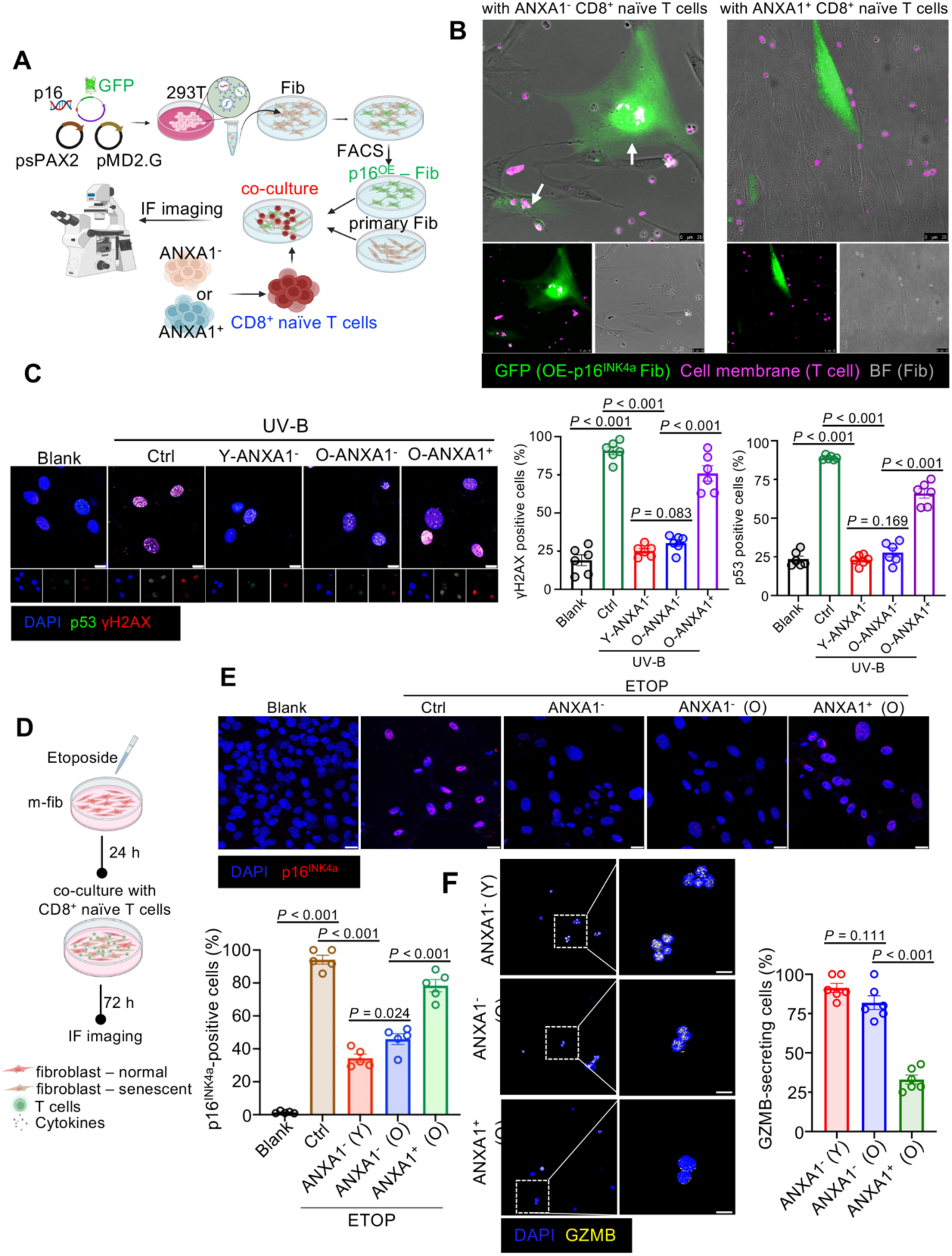
Senolytic capacity of ANXA1⁻ T cells is robust across multiple *in vitro* senescence models. **(A)** Schematic of the co-culture experiment using fibroblasts engineered to express p16^INK4a^-GFP as a model of senescent cells. Human ANXA1^−^ or ANXA1^+^ CD8^+^ naïve T cells using FACS were co-cultured with fibroblasts for live imaging. **(B)** Representative stills from live-cell imaging showing the targeting and clearance of p16^INK4a^-GFP⁺ senescent cells by ANXA1^−^ or ANXA1^+^ CD8^+^ naïve T cells 6 hours post-culture. **(C)** Representative immunofluorescence images and quantification of p53- and γH2AX- expressing fibroblasts after co-culture in UV-B induced senescence model (n = 6 per group), corresponding with Fig. 5B. **(D)** Schematic of an Etoposide-induced senescence model. **(E)** Representative immunofluorescence images and quantification showing a significant reduction of p16^INK4a^-positive senescent fibroblasts after co-culture with ANXA1⁻ T cells, but not with ANXA1⁺ T cells (n = 5 per group). Scale bar, 10 µm. **(F)** Representative immunofluorescence images and quantification of GZMB-secreting cells from different T cell groups after co-culture in (E) (n = 5 per group). Scale bar, 10 µm. Data are presented as mean ± SEM. *P*-values were determined by one-way ANOVA and two-tailed unpaired t-test.

**Fig. S8.**
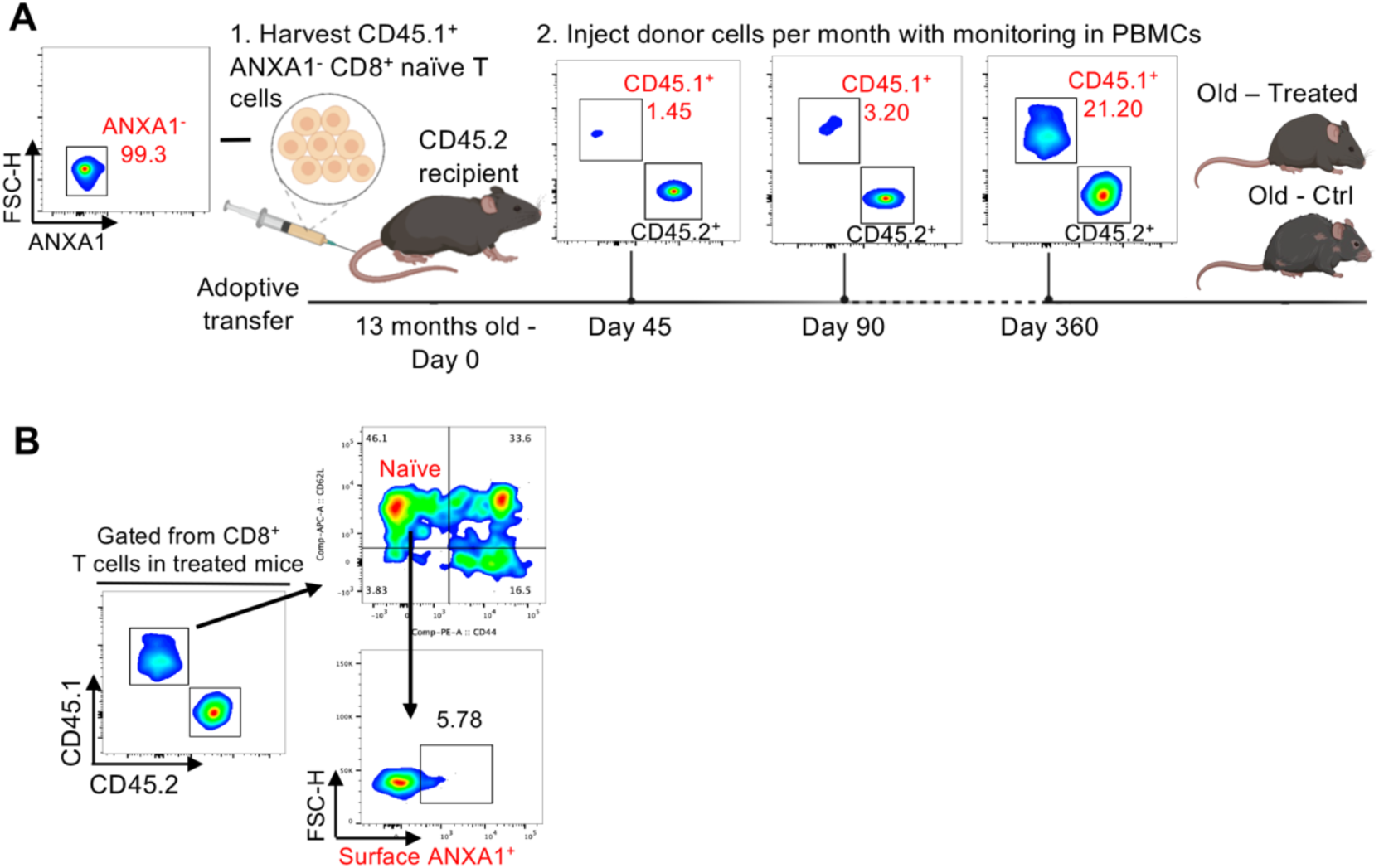
Adoptively transferred ANXA1⁻ CD8⁺ naïve T cells maintain a stable and functional phenotype in aged hosts. **(A)** Schematic diagram on adoptive transfer cell therapy model. CD45.1⁺ ANXA1⁻ CD8⁺ naïve T cells were adoptively transferred into aged CD45.2⁺ recipient mice, followed by monthly monitoring of donor cell presence and phenotype in peripheral blood. **(B)** Representative flow cytometry plots from peripheral blood at Day 360 post-transfer, showing the identification of CD45.1⁺ donor cells. Analysis of the donor cell population reveals that a significant fraction retains a naïve-like phenotype (CD62L⁺ CD44⁻) and remains ANXA1⁻.

**Fig. S9.**
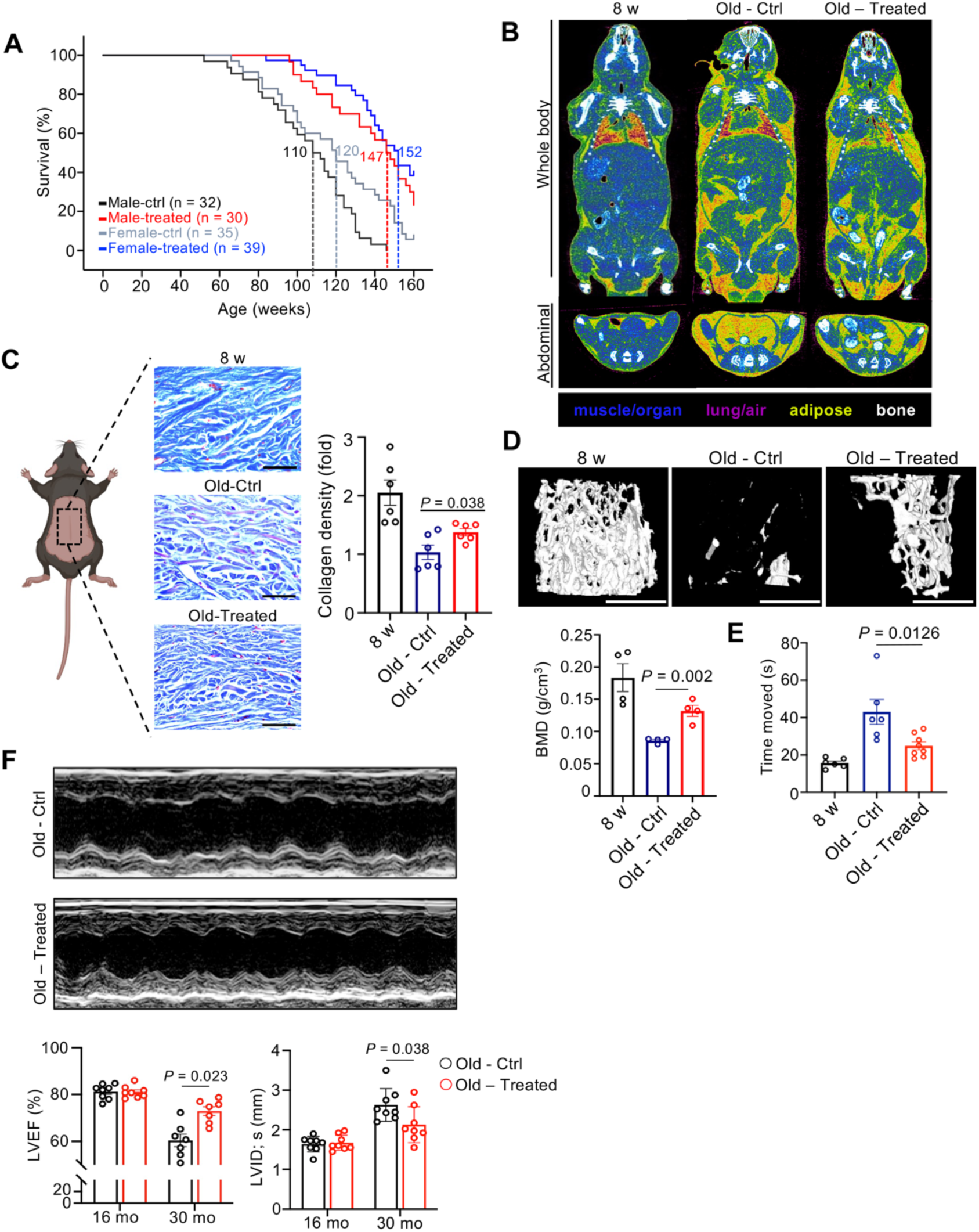
Cell therapy comprehensively improves multiple systemic healthspan indicators. **(A)** Kaplan-Meier survival curves for all mice split by genders. Statistical significance was determined by the Log-rank test. Median survival is indicated by dashed lines. **(B)** 3D micro-CT images of the whole body and abdominal area in 8 w (8-week-old), Old – Ctrl (30-month-old), and Old – Treated (30-month-old) mice. **(C)** Representative images and quantification of Masson’s staining on back skin of mice (n = 6 per group). Scale bar, 200 µm. **(D)** Representative micro-CT images and quantification of bone mineral density (BMD) of trabecular bone from different groups of mice (n = 4 per group). Scale bar = 1 mm. **(E)** Quantification of performance in the pole test for 3 groups of mice (n = 6 per group). **(F)** Representative M-mode echocardiographic images and corresponding measurements of ejection fraction (LVEF) and left ventricular end-systolic dimension (LVID; s) of 30-month-old mice in non-treated and treated group (n = 8 per group). Data are presented as mean ± SEM. *P*-values were determined by one-way ANOVA and two-tailed unpaired t-test.

**Fig. S10.**
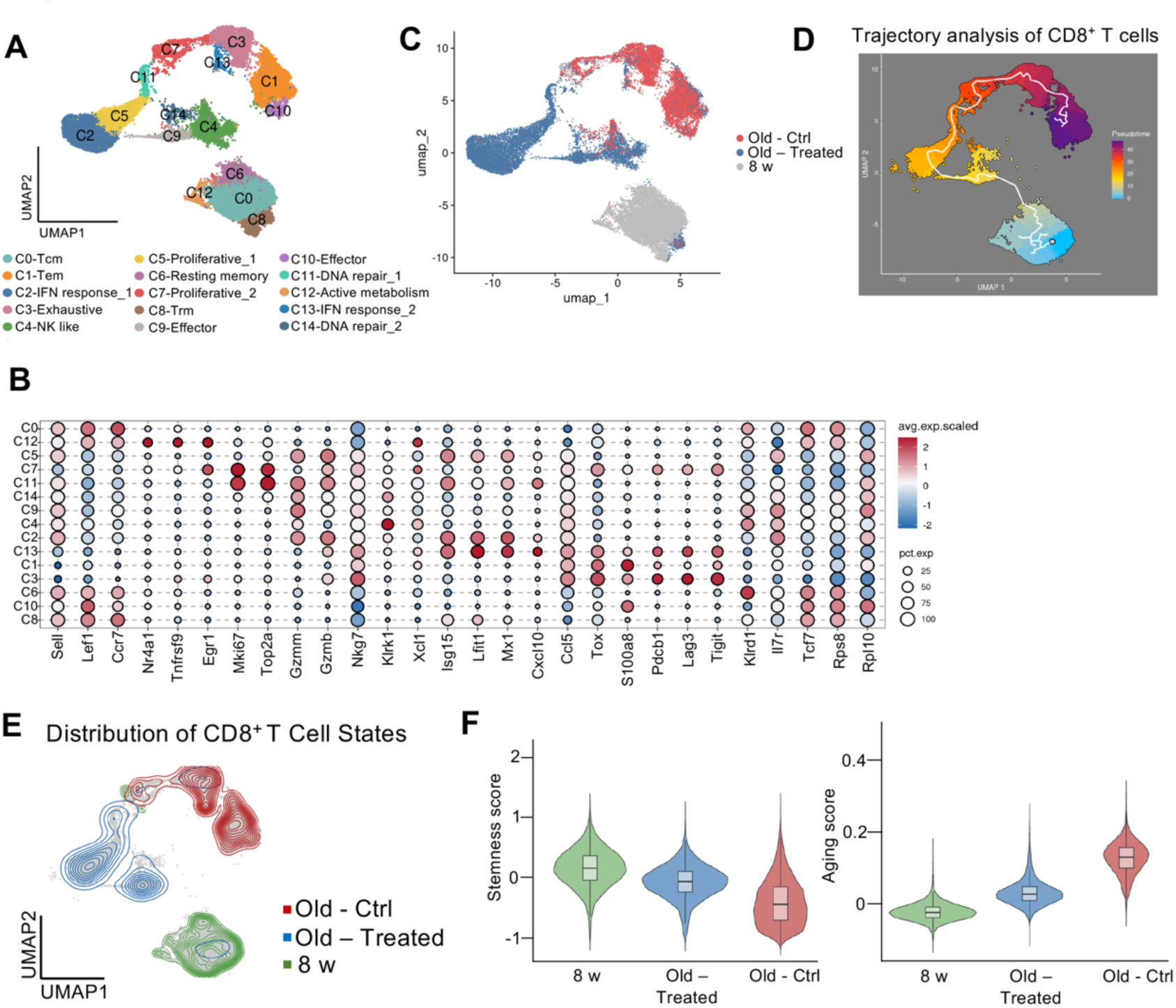
Therapeutic transfer of ANXA1⁻ T cells rejuvenates the endogenous CD8⁺ T cell compartment in the aged bone marrow. **(A)** UMAP visualization of re-clustered CD8⁺ T cells from the bone marrow of 8 w (8-week-old), Old-Ctrl, and Old-Treated mice, identifying 15 distinct subsets. Tcm, central memory T cell; Tem, effector memory T cell; Trm, tissue-resident memory T cell. **(B)** Dot plot showing the expression of key marker genes used to define the CD8⁺ T-cell subsets in (A). **(C)** UMAP projections of CD8⁺ T cells, split by the three experimental groups, illustrating a global shift in the cellular landscape after treatment. **(D)** Pseudotime trajectory analysis of all CD8⁺ T cells, ordered from a progenitor/memory state (blue/green) to an effector/dysfunctional state (purple). **(E)** Density contour plots showing the distribution of cells from each experimental group along the UMAP space defined in (**A**). Note the shift of the Old-Treated (blue) population away from the Old-Ctrl (red) distribution and towards the Young-ctrl (green) distribution. **(F)** Violin plots (**F**) comparing the “Stemness score” and “Aging score” for all CD8⁺ T cells from the three experimental groups.

**Table S1.**
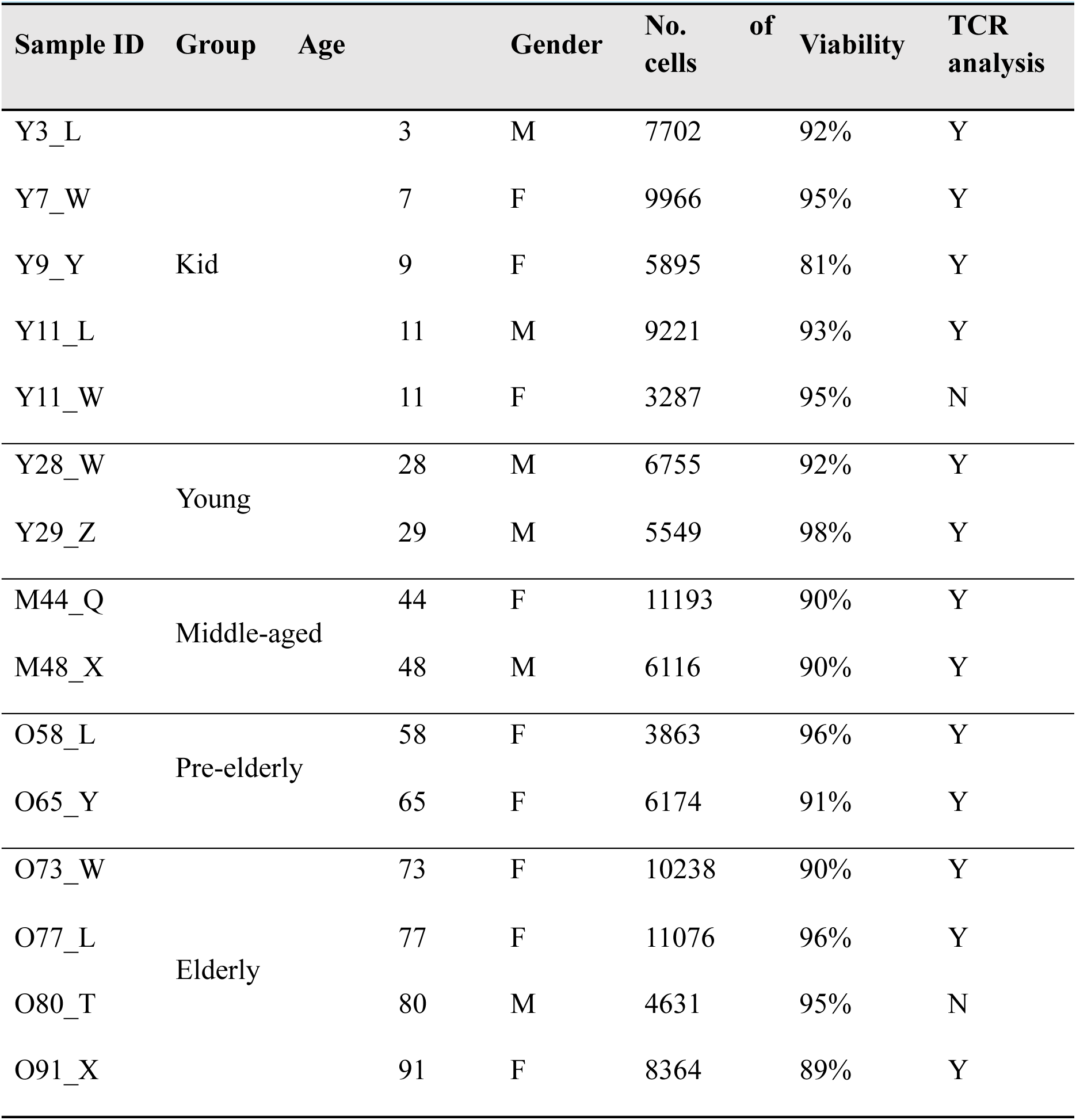
Sample information.

### Movie S1

Live-imaging video of ANXA1^−^ CD8^+^ naïve T cells with p16^INK4a^-GFP and primary fibroblasts.

