## supplementary figures for "Transplanting ANXA1⁻ CD8⁺ Naïve T cells Delay Aging Through Senolysis"

**Fig. S1**

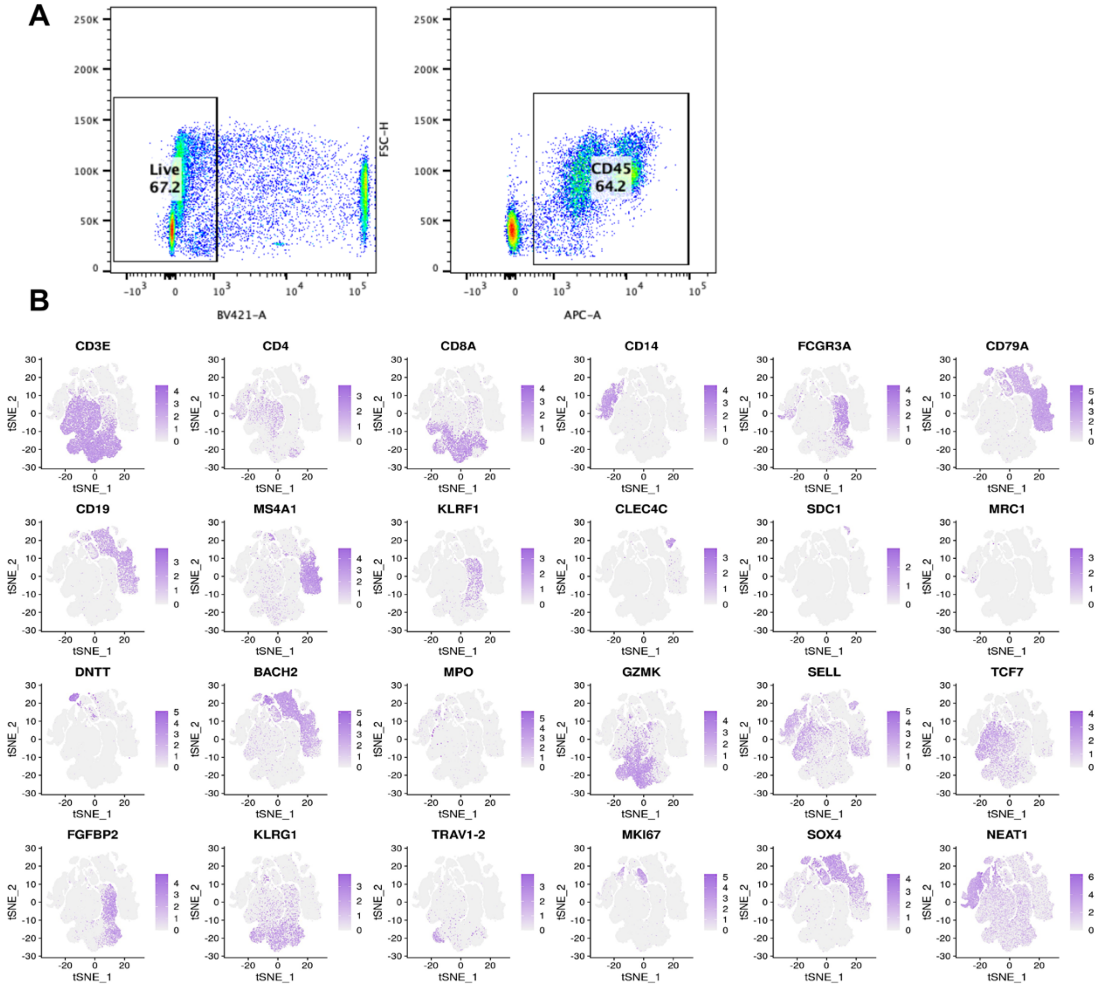


**Fig. S1. Quality control and cell lineage annotation of human bone marrow scRNA-seq data.**

(**A**) FACS gating strategy for isolating $CD45^+$ immune cells from human bone marrow for scRNA-seq and scTCR-seq analysis.

(**B**) Feature plots of canonical marker genes used to identify the major immune cell populations annotated in **Fig. 1B**.

**Fig. S2**

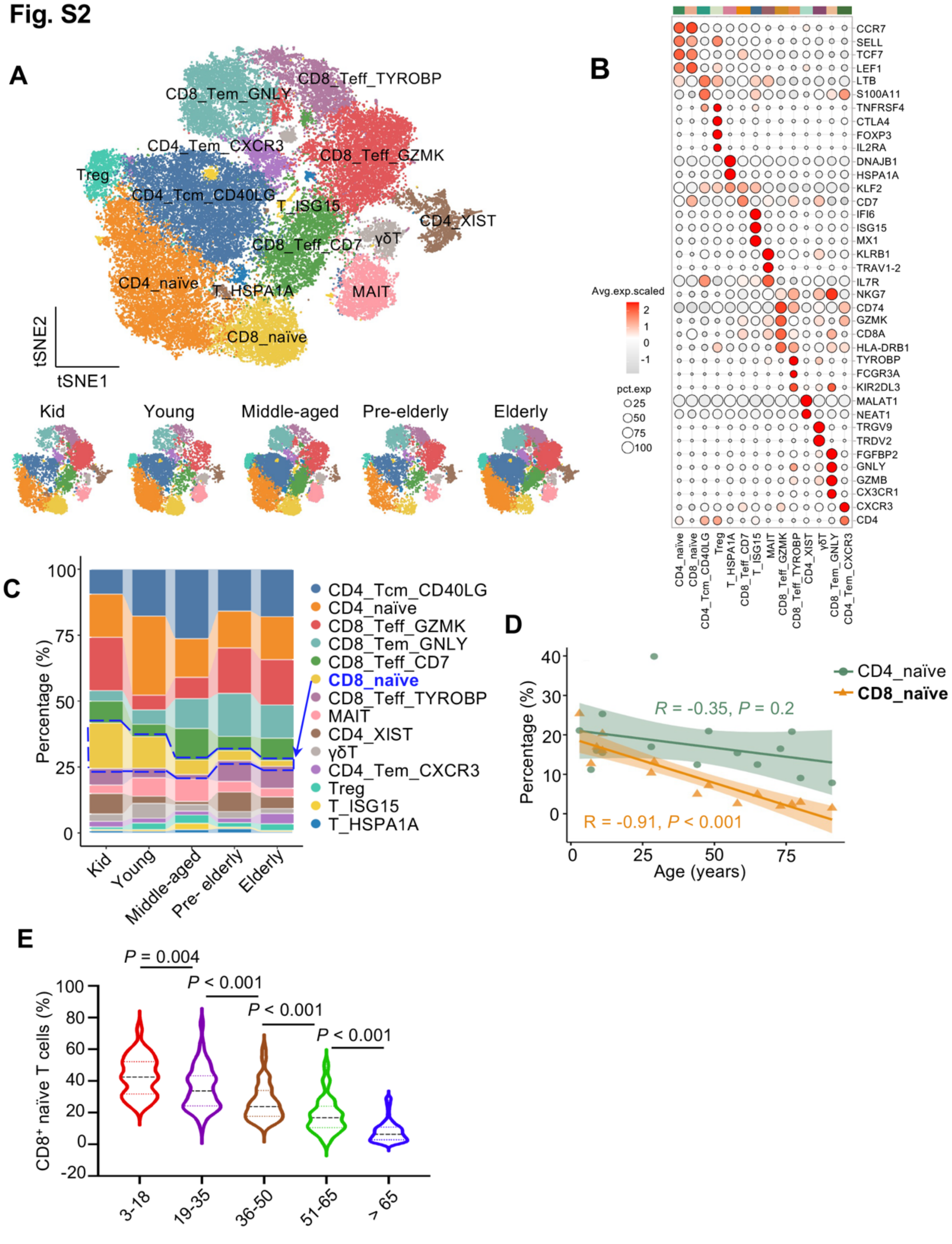
A
CD8_Tem_GNLY
CD8_Teff_TYROBP
CD4_Tem_CXCR3
CD8_Teff_GZMK
Treg
CD4_Tcm_CD40LG
T_ISG15
CD4_XIST
CD8_Teff_CD7
γδT
CD4_naïve
T_HSPA1A
MAIT
CD8_naïve
tSNE2
tSNE1
Kid
Young
Middle-aged
Pre-elderly
Elderly
B
Avg.exp.scaled
pct.exp
C
Percentage (%)
D
CD4_naïve
CD8_naïve
R = -0.35, P = 0.2
R = -0.91, P < 0.001
Percentage (%)
Age (years)
E
P = 0.004
P < 0.001
P < 0.001
P < 0.001
CD8+ naïve T cells (%)
3-18
19-35
36-50
51-65
> 65

**Fig. S2. scRNA-seq analysis of T cells in human bone marrow samples.**
(**A**) T-SNE clustering of all identified T-cell subsets from human bone marrow for scRNA-seq data. T cells from different groups of donors are shown separately.

(**B**) Dot plot on signature genes identified in each T cell subsets. Color scale represents the average expression level of selected genes and dot size represents the proportion of cells expressing marker genes in each cluster. Names of each cell type are listed below, corresponding with those in (**A**)**.**

(**C**) Stacked bar plot showing the proportional changes of cell clusters between 5 age groups.

(**D**) Correlation between the proportion of naïve T-cell subsets and donor age. Scatter plots show the percentage of $CD4^+$ and $CD8^+$ naïve T cells relative to total T cells versus donor age. Solid lines indicate linear regression fits, with Pearson correlation coefficient (R) and *P*-values displayed.

(**E**) Validation of age-related decline in $CD8^+$ naïve T-cell frequency in PBMCs. Box plot showing the percentage of $CD8^+$ naïve T cells (identified by flow cytometry as in Fig. 2E) within total $CD8^+$ T cells across age groups in the large validation cohort (n=388, same cohort as Fig. 2F).

**Fig. S3**

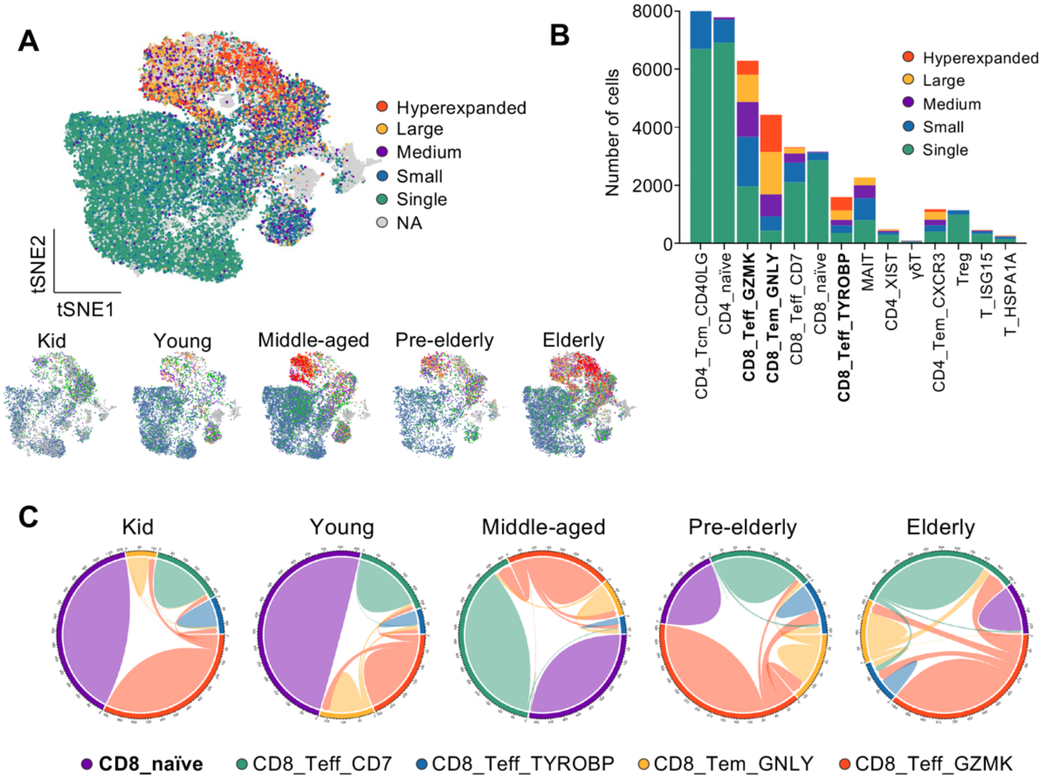


**D**

Hyperexpanded TCR

| CDR3-TRA | CDR3-TRB | Frequency | Cluster |
|---|---|---|---|
| CAHNTGNQFYF | CASSPITGTGSYGYTF | 263 | CD8_Teff_GZMK |
| CAVGTSYDKVIF | CASSVPGPDNEQFF | 241 | CD8_Tem_GNLY |
| CAMREEPFGNEKLTF | CASSRGLAGRQETQYF | 196 | CD8_Teff_TYROBP |

**Fig. S3. scTCR-seq analysis of the T-cell receptor repertoire.**

(**A**) T cell receptor (TCR) clone-types analysis in scTCR-seq (n=13) projected on t-SNE plot in Fig. S2A. Colors represent different clonotypes based on numbers (x) of same clonotype. 6 levels are defined, including: hyperexpanded ($100 < x \leq 500$), large ($20 < x \leq 100$), medium ($5 < x \leq 20$), small ($1 < x \leq 5$), single ($x = 1$) and NA. Separated clustering plots are shown below from samples of different age groups.

(**B**) Distribution of 5 clonotypes in all cell clusters. Bold texts represent clusters with different levels of clonotypes, especially with larger expanded clones

(**C**) Chord diagram shows the repertoire changes in $CD8^+$ T cells compared between samples in different age groups.

(**D**) Top 3 hyperexpanded TCR-CDR3 sequences are listed, all enriched in $CD8^+$ T cells.

**Fig. S4**

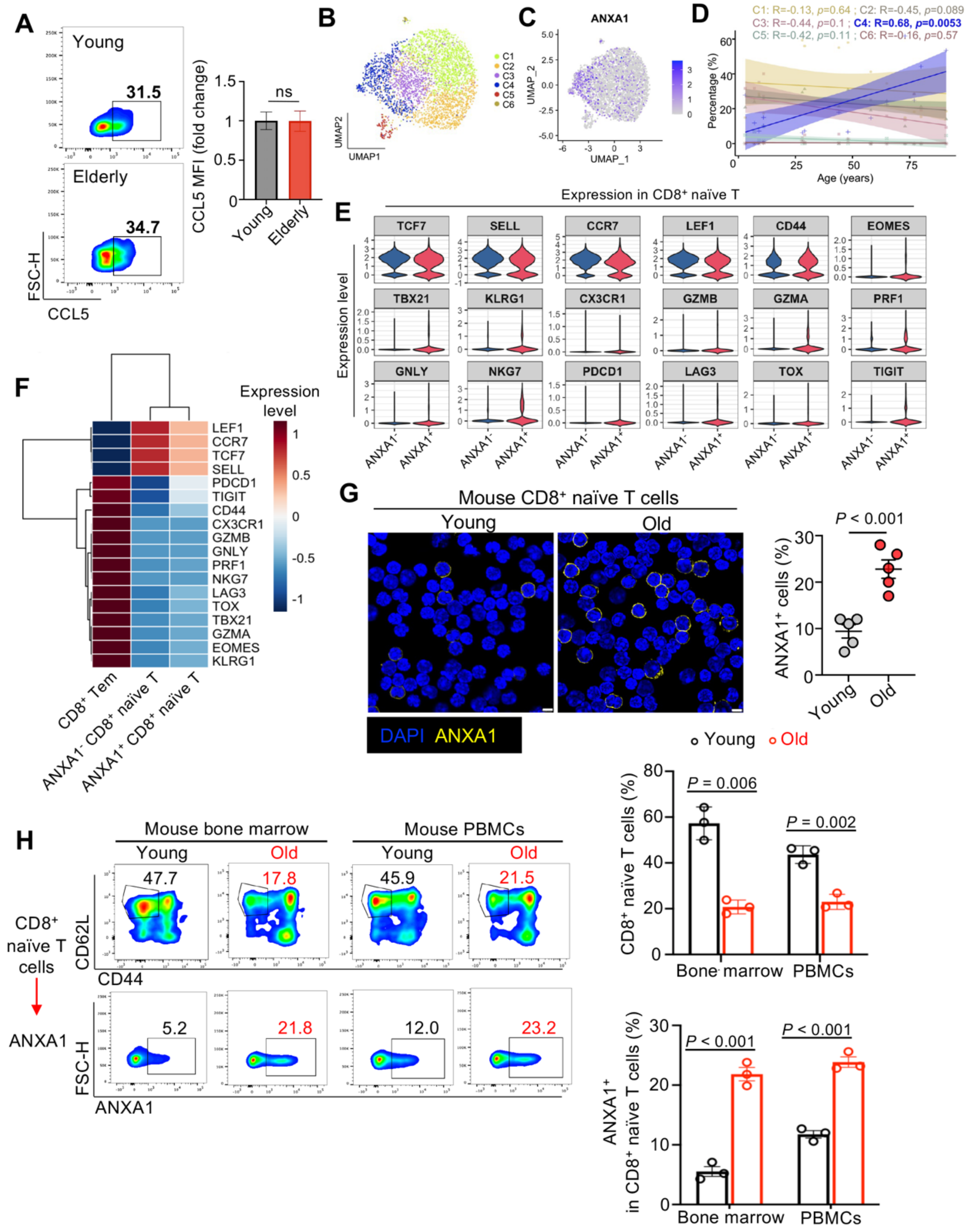
A
Young
31.5
Elderly
34.7
FSC-H
CCL5
CCL5 MFI (fold change)
ns
Young
Elderly
B
UMAP1
UMAP2
C1
C2
C3
C4
C5
C6
C
ANXA1
UMAP_1
UMAP_2
D
C1: R=-0.13, p=0.64 ; C2: R=-0.45, p=0.089
C3: R=-0.44, p=0.1 ; C4: R=0.68, p=0.0053
C5: R=-0.42, p=0.11 ; C6: R=-0.16, p=0.57
Percentage (%)
Age (years)
E
Expression in CD8⁺ naïve T
Expression level
TCF7
SELL
CCR7
LEF1
CD44
EOMES
TBX21
KLRG1
CX3CR1
GZMB
GZMA
PRF1
GNLY
NKG7
PDCD1
LAG3
TOX
TIGIT
ANXA1⁻
ANXA1⁺
F
Expression level
LEF1
CCR7
TCF7
SELL
PDCD1
TIGIT
CD44
CX3CR1
GZMB
GNLY
PRF1
NKG7
LAG3
TOX
TBX21
GZMA
EOMES
KLRG1
CD8⁺ Tem
ANXA1⁻ CD8⁺ naïve T
ANXA1⁺ CD8⁺ naïve T
G
Mouse CD8⁺ naïve T cells
Young
Old
DAPI ANXA1
P < 0.001
ANXA1⁺ cells (%)
Young
Old
H
Mouse bone marrow
Mouse PBMCs
Young
Old
47.7
17.8
45.9
21.5
CD8⁺ naïve T cells
ANXA1
CD62L
CD44
5.2
21.8
12.0
23.2
FSC-H
ANXA1
Young
Old
P = 0.006
P = 0.002
CD8⁺ naïve T cells (%)
Bone marrow
PBMCs
P < 0.001
P < 0.001
ANXA1⁺ in CD8⁺ naïve T cells (%)
Bone marrow
PBMCs

**Fig. S4. Phenotypic definition and validations of ANXA1$^-$ and ANXA1$^+$ CD8$^+$ T cells.**

(**A**) Representative flow cytometric analysis of CCL5 expression on CD8$^+$ naïve T cells in samples of young (19-35 y) and old (>65 y) group (n= 5 per group).

(**B**) Sub-clustering of CD8$^+$ naïve T cells from the scRNA-seq dataset. UMAP projection showing distinct subclusters (C1-C6) identified within the CD8$^+$ naïve T cell population.

(**C**) Feature plot overlaying normalized ANXA1 expression onto the UMAP projection from (**B**), highlighting its specific enrichment in the C4 subcluster.

(**D**) Scatter plots showing the correlation between the relative abundance of each subcluster (as a percentage of total CD8$^+$ naïve T cells) and donor age. Lines represent linear regression fits; Pearson correlation coefficient (R) and *P*-values are indicated.

(**E** and **F**) Violin plots (**E**) and heatmap (**F**) validating the naïve identity of both ANXA1$^-$ and ANXA1$^+$ CD8$^+$ naïve T cells defined in human scRNA-seq data. Average expression of canonical lineage markers for naïve, memory, effector, and exhausted T cells is compared across three populations with CD8$^+$ effector memory (Tem) cells as a non-naïve control. Expression values are scaled per row.

(**G**) Representative immunofluorescence images and quantification of ANXA1 surface expression on mouse CD8$^+$ naïve T cells from Young (8-week-old, n = 5) and Old (13-month-old, n = 5) mice. Scale bar, 5 μm.

(**H**) Representative flow cytometry results and quantifications on the percentage of CD8$^+$ naïve T cells and surface ANXA1 expression in young and old mice (n = 3 per group).

Data in quantification plots are presented as mean ± SEM. Statistical significance was determined by one-way ANOVA and two-tailed unpaired t-test.

Fig. S5

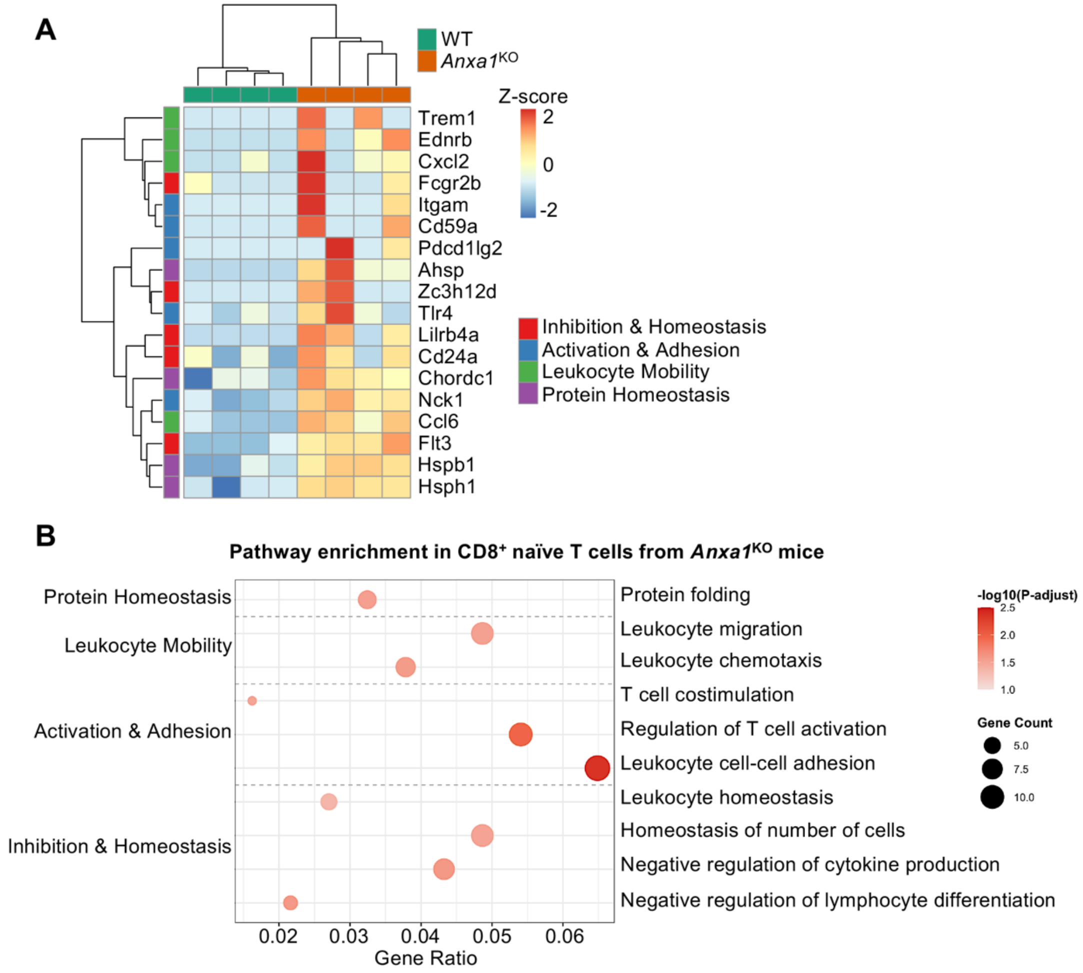


**Fig. S5. ANXA1-deficient CD8$^+$ naïve T cells exhibit a primed phenotype at basal state.**

(**A**) Heatmap of DEGs from RNA-seq of resting WT and *Anxa1*$^{KO}$ CD8$^+$ naïve T cells.

(**B**) GO enrichment bubble plot of pathways upregulated in *Anxa1*$^{KO}$ versus WT CD8$^+$ naïve T cells at basal state.

**Fig. S6**

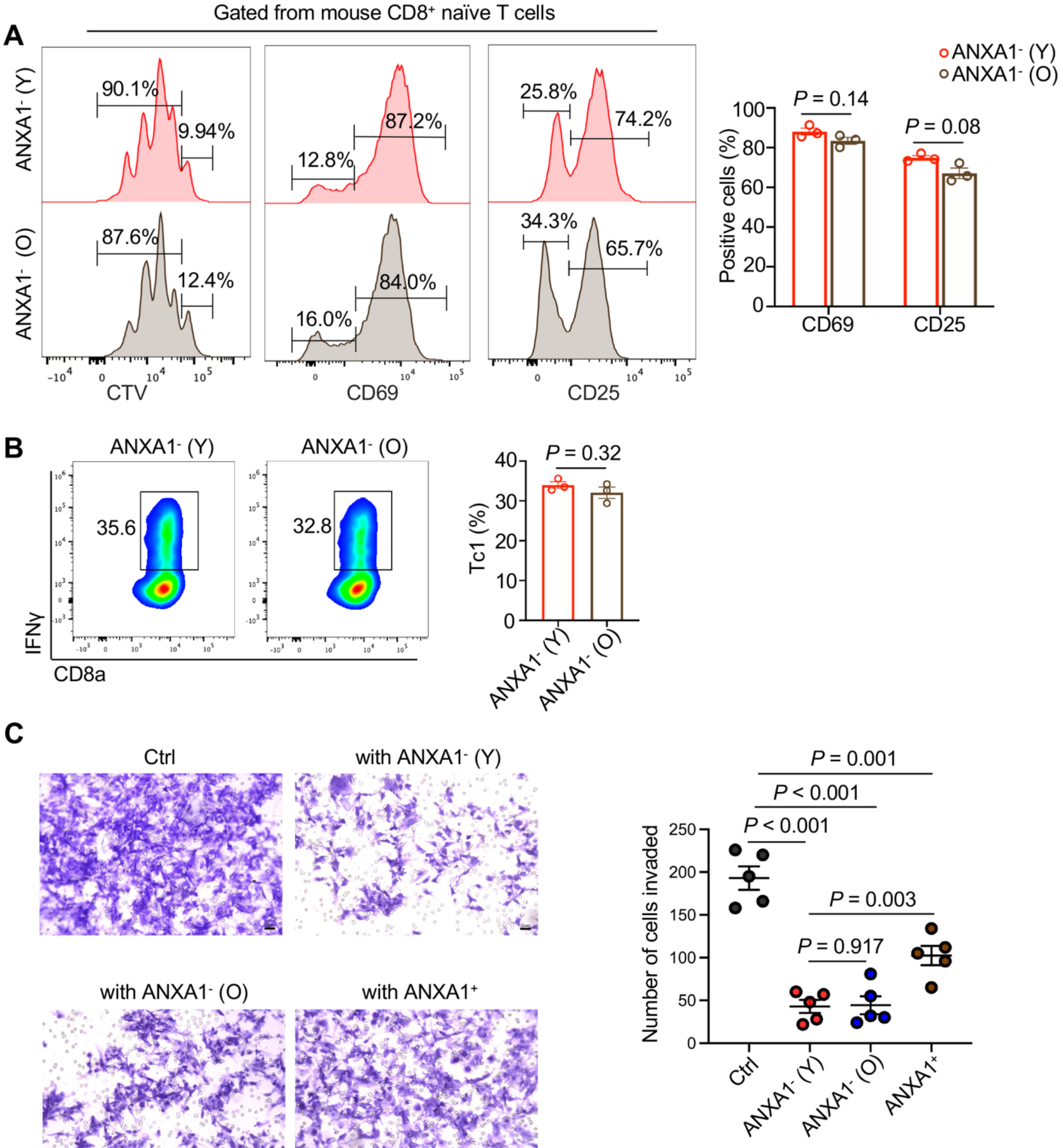
A
Gated from mouse CD8+ naïve T cells
ANXA1- (Y)
ANXA1- (O)
90.1%
9.94%
87.6%
12.4%
12.8%
87.2%
16.0%
84.0%
25.8%
74.2%
34.3%
65.7%
CTV
CD69
CD25
ANXA1- (Y)
ANXA1- (O)
Positive cells (%)
P = 0.14
P = 0.08
CD69
CD25
B
ANXA1- (Y)
ANXA1- (O)
35.6
32.8
IFNγ
CD8a
Tc1 (%)
P = 0.32
ANXA1- (Y)
ANXA1- (O)
C
Ctrl
with ANXA1- (Y)
with ANXA1- (O)
with ANXA1+
Number of cells invaded
P = 0.001
P < 0.001
P < 0.001
P = 0.003
P = 0.917
Ctrl
ANXA1- (Y)
ANXA1- (O)
ANXA1+

**Fig. S6. ANXA1$^-$ CD8$^+$ naïve T cells cells from young and old mice are functionally comparable *in vitro*.**

(**A**) Representative flow cytometry histograms and quantifications of CTV-staining, CD69 and CD25 expression of ANXA1$^-$ CD8$^+$ naïve T cells from young and aged mice (n = 3 per group).

(**B**) Representative flow cytometry results and quantifications of Tc1 differentiation of ANXA1$^-$ CD8$^+$ naïve T cells from young and aged mice.

(**C**) Representative images and quantification from a transwell invasion assay showing that both young and old mouse ANXA1$^-$ T cells, but not ANXA1$^+$ T cells (n = 5 per group), significantly inhibit the invasion of U-2 OS tumor cells, with arrows indicate tumor cells passed the transwell membrane.

Data are presented as mean ± SEM. *P*-values were determined by one-way ANOVA and two-tailed unpaired t-test.

**Fig. S7**

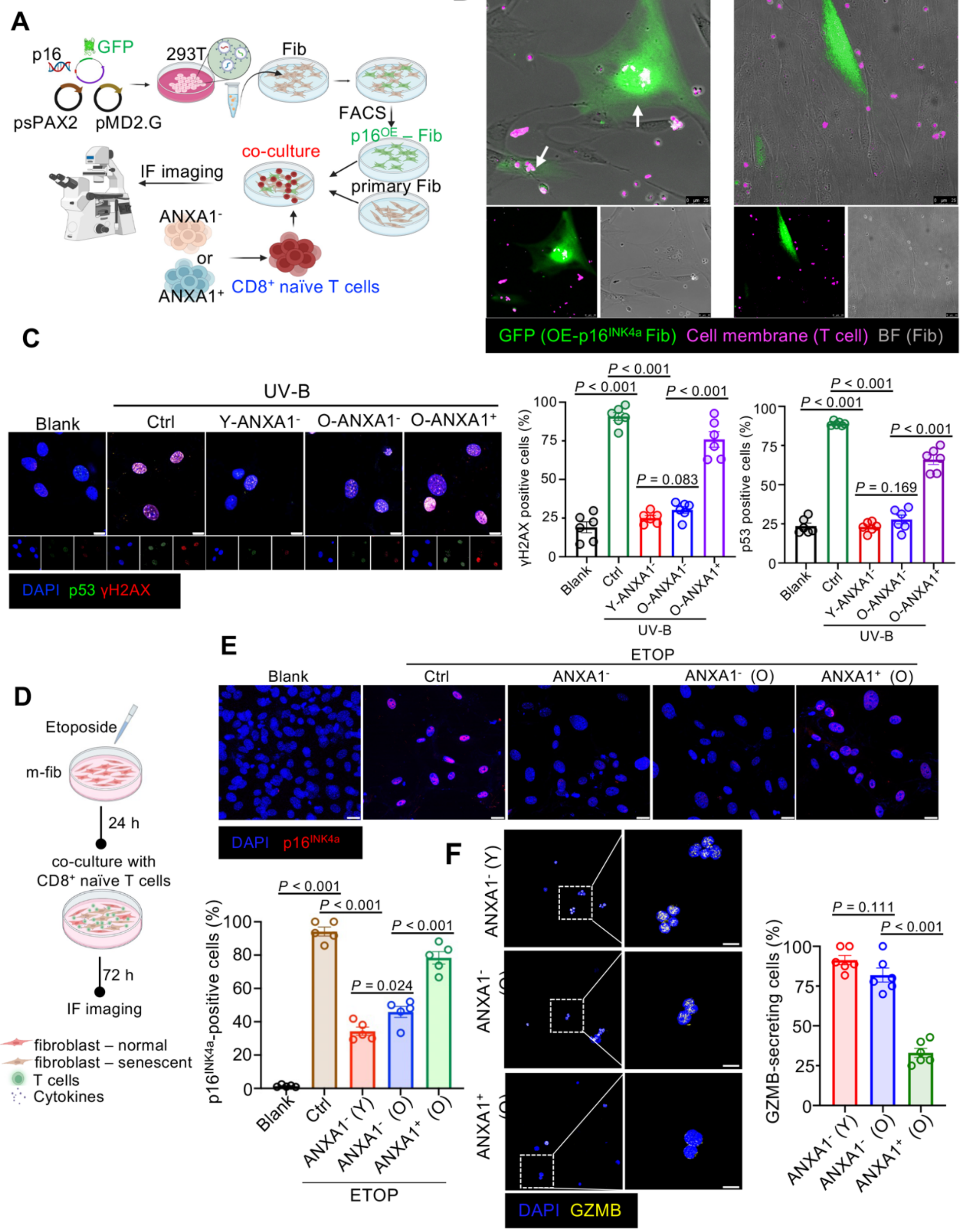
A
p16
GFP
293T
Fib
psPAX2
pMD2.G
FACS
co-culture
IF imaging
primary Fib
ANXA1-
or
ANXA1+
CD8+ naïve T cells
B
with ANXA1- CD8+ naïve T cells
with ANXA1+ CD8+ naïve T cells
GFP (OE-p16INK4a Fib) Cell membrane (T cell) BF (Fib)
C
UV-B
Blank
Ctrl
Y-ANXA1-
O-ANXA1-
O-ANXA1+
DAPI p53 γH2AX
γH2AX positive cells (%)
p53 positive cells (%)
P < 0.001
P = 0.083
P = 0.169
D
Etoposide
m-fib
24 h
co-culture with
CD8+ naïve T cells
72 h
IF imaging
fibroblast – normal
fibroblast – senescent
T cells
Cytokines
E
ETOP
ANXA1- (O)
ANXA1+ (O)
DAPI p16INK4a
p16INK4a-positive cells (%)
P = 0.024
F
GZMB-secreting cells (%)
P = 0.111
DAPI GZMB

**Fig. S7. Senolytic capacity of ANXA1⁻ T cells is robust across multiple *in vitro* senescence models.**

(**A**) Schematic of the co-culture experiment using fibroblasts engineered to express $p16^{INK4a}$-GFP as a model of senescent cells. Human $ANXA1^-$ or $ANXA1^+$ $CD8^+$ naïve T cells using FACS were co-cultured with fibroblasts for live imaging.

(**B**) Representative stills from live-cell imaging showing the targeting and clearance of $p16^{INK4a}$-$GFP^+$ senescent cells by $ANXA1^-$ or $ANXA1^+$ $CD8^+$ naïve T cells 6 hours post-culture.

(**C**) Representative immunofluorescence images and quantification of p53- and γH2AX-expressing fibroblasts after co-culture in UV-B induced senescence model (n = 6 per group), corresponding with Fig. 5B.

(**D**) Schematic of an Etoposide-induced senescence model.

(**E)** Representative immunofluorescence images and quantification showing a significant reduction of $p16^{INK4a}$-positive senescent fibroblasts after co-culture with $ANXA1^-$ T cells, but not with $ANXA1^+$ T cells (n = 5 per group). Scale bar, 10 μm.

(**F**) Representative immunofluorescence images and quantification of GZMB-secreting cells from different T cell groups after co-culture in (E) (n = 5 per group). Scale bar, 10 μm.

Data are presented as mean ± SEM. *P*-values were determined by one-way ANOVA and two-tailed unpaired t-test.

**Fig. S8**

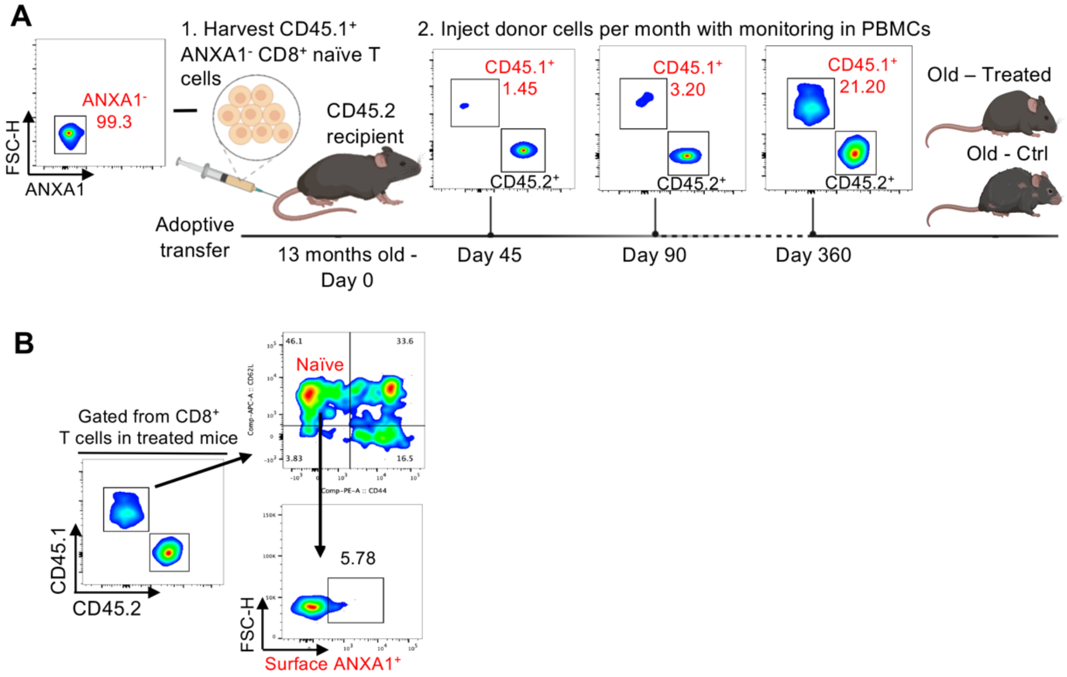


**Fig. S8. Adoptively transferred ANXA1⁻ CD8⁺ naïve T cells maintain a stable and functional phenotype in aged hosts.**

(**A**) Schematic diagram on adoptive transfer cell therapy model. CD45.1⁺ ANXA1⁻ CD8⁺ naïve T cells were adoptively transferred into aged CD45.2⁺ recipient mice, followed by monthly monitoring of donor cell presence and phenotype in peripheral blood.

(**B**) Representative flow cytometry plots from peripheral blood at Day 360 post-transfer, showing the identification of CD45.1⁺ donor cells. Analysis of the donor cell population reveals that a significant fraction retains a naïve-like phenotype (CD62L⁺ CD44⁻) and remains ANXA1⁻.

Fig. S9

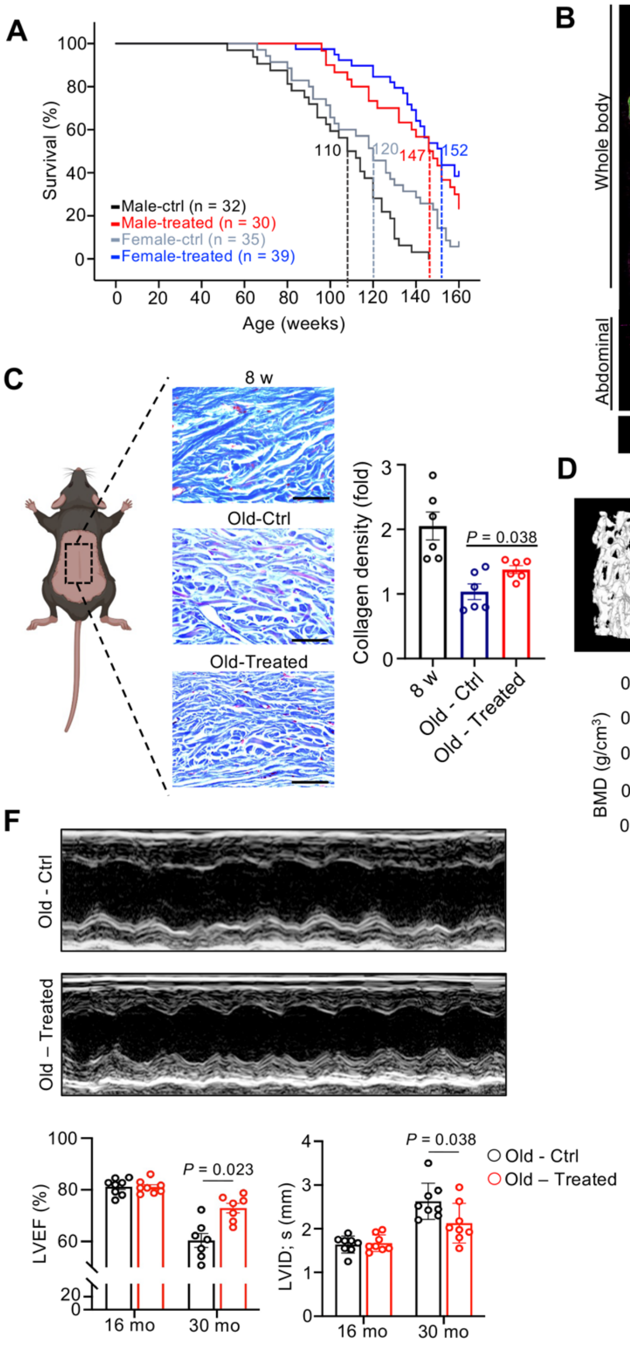
A
Survival (%)
Age (weeks)
110
120
147
152
Male-ctrl (n = 32)
Male-treated (n = 30)
Female-ctrl (n = 35)
Female-treated (n = 39)
B
8 w
Old - Ctrl
Old – Treated
Whole body
Abdominal
muscle/organ
lung/air
adipose
bone
C
8 w
Old-Ctrl
Old-Treated
Collagen density (fold)
P = 0.038
D
8 w
Old - Ctrl
Old – Treated
BMD (g/cm3)
P = 0.002
E
Time moved (s)
P = 0.0126
F
Old - Ctrl
Old – Treated
LVEF (%)
P = 0.023
16 mo
30 mo
LVID; s (mm)
P = 0.038
Old - Ctrl
Old – Treated

**Fig. S9. Cell therapy comprehensively improves multiple systemic healthspan indicators.**
(**A**) Kaplan-Meier survival curves for all mice split by genders. Statistical significance was determined by the Log-rank test. Median survival is indicated by dashed lines.

(**B**) 3D micro-CT images of the whole body and abdominal area in 8 w (8-week-old), Old – Ctrl (30-month-old), and Old – Treated (30-month-old) mice.

(**C**) Representative images and quantification of Masson's staining on back skin of mice (n = 6 per group). Scale bar, 200 μm.

(**D**) Representative micro-CT images and quantification of bone mineral density (BMD) of trabecular bone from different groups of mice (n = 4 per group). Scale bar = 1 mm.

(**E**) Quantification of performance in the pole test for 3 groups of mice (n = 6 per group).

(**F**) Representative M-mode echocardiographic images and corresponding measurements of ejection fraction (LVEF) and left ventricular end-systolic dimension (LVID; s) of 30-month-old mice in non-treated and treated group (n = 8 per group).

Data are presented as mean ± SEM. *P*-values were determined by one-way ANOVA and two-tailed unpaired t-test.

### Fig. S10

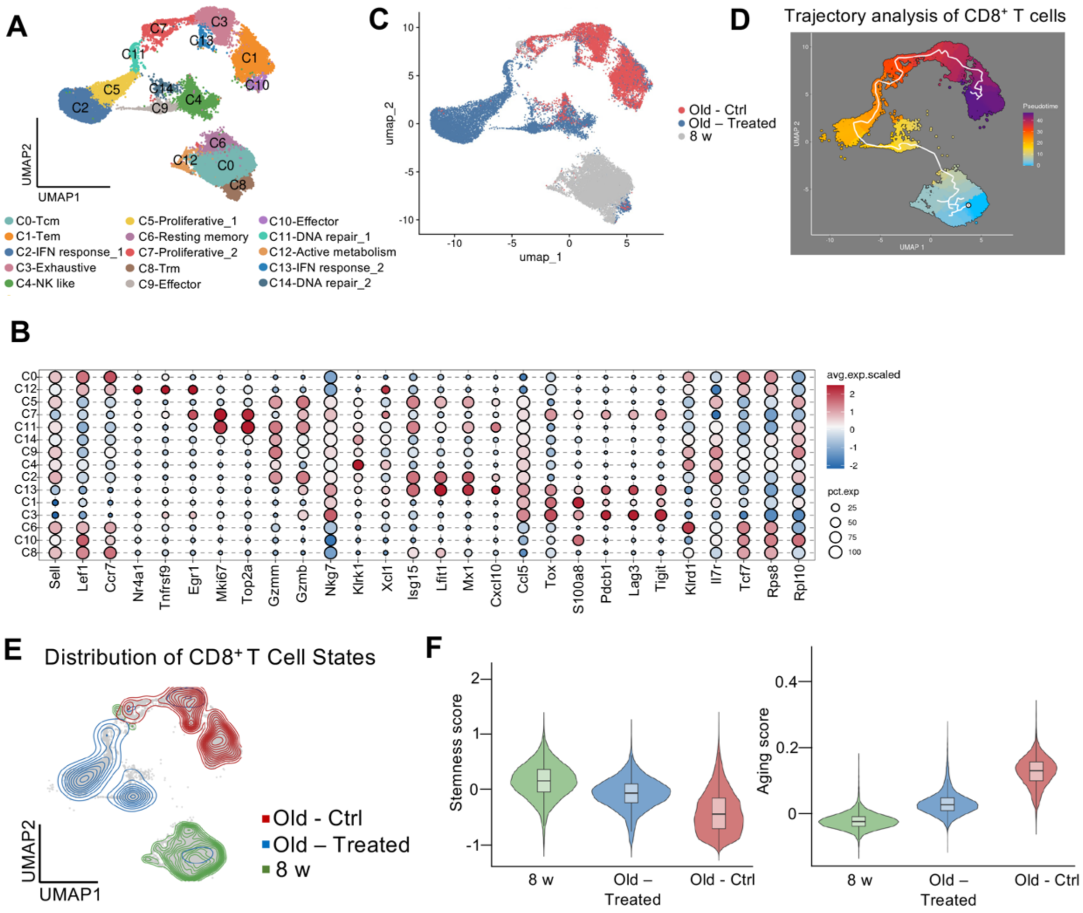
A
C7
C3
C13
C11
C1
C10
C5
C14
C2
C9
C4
C6
C12
C0
C8
UMAP2
UMAP1
C0-Tcm
C1-Tem
C2-IFN response_1
C3-Exhaustive
C4-NK like
C5-Proliferative_1
C6-Resting memory
C7-Proliferative_2
C8-Trm
C9-Effector
C10-Effector
C11-DNA repair_1
C12-Active metabolism
C13-IFN response_2
C14-DNA repair_2
C
umap_2
umap_1
Old - Ctrl
Old – Treated
8 w
D
Trajectory analysis of CD8⁺ T cells
Pseudotime
B
avg.exp.scaled
pct.exp
25
50
75
100
Sell
Lef1
Ccr7
Nr4a1
Tnfrsf9
Egr1
Mki67
Top2a
Gzmm
Gzmb
Nkg7
Klrk1
Xcl1
Isg15
Ifit1
Mx1
Cxcl10
Ccl5
Tox
S100a8
Pdcb1
Lag3
Tigit
Klrd1
Il7r
Tcf7
Rps8
Rpl10
E
Distribution of CD8⁺ T Cell States
UMAP2
UMAP1
Old - Ctrl
Old – Treated
8 w
F
Stemness score
Aging score
8 w
Old – Treated
Old - Ctrl

**Fig. S10. Therapeutic transfer of ANXA1$^{-}$ T cells rejuvenates the endogenous CD8$^{+}$ T cell compartment in the aged bone marrow.**

(**A**) UMAP visualization of re-clustered CD8$^{+}$ T cells from the bone marrow of 8 w (8-week-old), Old-Ctrl, and Old-Treated mice, identifying 15 distinct subsets. Tcm, central memory T cell; Tem, effector memory T cell; Trm, tissue-resident memory T cell.

(**B**) Dot plot showing the expression of key marker genes used to define the CD8$^{+}$ T-cell subsets in (A).

(**C**) UMAP projections of CD8$^{+}$ T cells, split by the three experimental groups, illustrating a global shift in the cellular landscape after treatment.

(**D**) Pseudotime trajectory analysis of all CD8$^{+}$ T cells, ordered from a progenitor/memory state (blue/green) to an effector/dysfunctional state (purple).

(**E**) Density contour plots showing the distribution of cells from each experimental group along the UMAP space defined in (**A**). Note the shift of the Old-Treated (blue) population away from the Old-Ctrl (red) distribution and towards the Young-ctrl (green) distribution.

(**F**) Violin plots (**F**) comparing the "Stemness score" and "Aging score" for all CD8$^{+}$ T cells from the three experimental groups.

| Table S1. Sample information | | | | | | |
|---|---|---|---|---|---|---|
| **Sample ID** | **Group** | **Age** | **Gender** | **No. of cells** | **Viability** | **TCR analysis** |
| Y3_L | Kid | 3 | M | 7702 | 92% | Y |
| Y7_W | | 7 | F | 9966 | 95% | Y |
| Y9_Y | | 9 | F | 5895 | 81% | Y |
| Y11_L | | 11 | M | 9221 | 93% | Y |
| Y11_W | | 11 | F | 3287 | 95% | N |
| Y28_W | Young | 28 | M | 6755 | 92% | Y |
| Y29_Z | | 29 | M | 5549 | 98% | Y |
| M44_Q | Middle-aged | 44 | F | 11193 | 90% | Y |
| M48_X | | 48 | M | 6116 | 90% | Y |
| O58_L | Pre-elderly | 58 | F | 3863 | 96% | Y |
| O65_Y | | 65 | F | 6174 | 91% | Y |
| O73_W | Elderly | 73 | F | 10238 | 90% | Y |
| O77_L | | 77 | F | 11076 | 96% | Y |
| O80_T | | 80 | M | 4631 | 95% | N |
| O91_X | | 91 | F | 8364 | 89% | Y |

**Movie S1.**

Live-imaging video of ANXA1$^{-}$ CD8$^{+}$ naïve T cells with p16$^{INK4a}$-GFP and primary fibroblasts.
